# High prevalence of antibiotic resistance in traditionally fermented foods as a critical risk factor for host gut antibiotic resistome

**DOI:** 10.1101/2023.04.21.537834

**Authors:** Yutong Li, Siying Fu, Matthias S. Klein, Hua Wang

**Affiliations:** Department of Food Science and Technology, The Ohio State University. 2015 Fyffe Court, Columbus, OH 43210, USA

**Keywords:** Gut microbiota, traditionally fermented foods, intervention, antibiotic resistance, resistome, opportunistic pathogens, kimchi, artisan cheese

## Abstract

Disrupted gut microbiota as a critical risk factor for many noncommunicable diseases is largely driven by gut microbiota-impacting drugs, especially orally administrated as well as biliary excreted antibiotics. Fermented food consumption has been encouraged to replenish disrupted gut microbiota, but its overall impact on host gut health remains to be elucidated. This study examined retail traditionally fermented foods and gut microbiota of consumers of fermented foods for antibiotic resistome. Dietary intervention by fermented foods was found leading to a surge of the antibiotic resistome in gut microbiota of most human subjects. Antibiotic resistome was further illustrated in traditionally fermented food samples, and viable antibiotic resistant (AR) bacteria were recovered and highly prevalent in retail kimchi and artisan cheeses assessed in this pilot screening. Identified AR isolates included pathogens of importance in nosocomial infections such as *Klebsiella pneumoniae*, *Enterococcus*, etc., as well as commensals and lactic acid bacteria, some exhibited extremely high minimum inhibitory concentration (MIC) against antibiotics of clinical significance. Exposing fermented food microbiota to representative antibiotics further led to a boost of the corresponding antibiotic and multidrug-resistance gene pools and disturbed microbiota. These results revealed an underestimated public health risk associated with fermented foods intervention, particularly to susceptible population with gastrointestinal tract symptoms and compromised immune functions seeking gut microbiota rescue. The findings call for more comprehensive investigation and investment on the benefits and potential safety challenges associated with traditionally fermented foods, productive intervention of foodborne antibiotic resistance, and strategic movements to mitigate unnecessary damages to the host gut microbiota.

## Introduction

The rapid rise of antibiotic resistance (AR) negates effective treatment of bacterial infections, shaking the foundation of modern medicine. AR bacteria in host gastrointestinal tract further indirectly contribute to gut microbiota dysbiosis resulting from even short-term antibiotic treatment (Zhou et al., 2020). Gut microbiota destruction has been recognized as a critical, shared risk factor for a growing list of noncommunicable “modern” diseases, ranging from malfunction of the host immune system, type-II diabetes, brain/neurological disorders, *Clostridium difficile* infections, to certain cardiovascular diseases and cancers (Sekirov et al., 2010; Chung et al., 2012; Foster et al., 2013; Blaser, 2016; Lynch & Pedersen, 2016; Smits et al., 2016; Meng et al., 2018; Gurung et al., 2020; Witkowski et al., 2020). Both AR and noncommunicable diseases are among the top global public health threats in the 21^st^ century (World Health Organization (WHO), 2019).

For decades, the broad applications of antibiotics have been blamed for the rapid surge of AR, and more recently, for disrupted host gut microbiota and associated “modern” diseases. But despite limiting the uses of antibiotics has been the primary control strategy worldwide, even essential antibiotic applications for infection prevention and treatment still cause irreversible damages in host gut microbiota (Zimmermann & Curtis, 2019; Wang, 2022). AR is a complicated issue with multiple risk factors (Wang, 2009; Wang et al., 2019). Particularly, gut impacting antibiotics, i.e., the mainstream oral administration of antibiotics, and using drugs with primary biliary instead of renal excretion, rather than the application of antibiotics, have been the key and shared driver for the rapid surge of the antibiotic resistance gene (ARG) pool, massively disrupted gut microbiota, and the rise of pathogens in gut microbiota (Zhang et al., 2013; Zhou et al., 2020). This paradigm-changing discovery is further supported by clinical observations on the trends of AR worldwide (Luber et al., 1996; Zhou et al., 2020). The finding is also applicable to non-antibiotic drugs with impact on gut microbiota (Weersma et al., 2020). According to CDC, over 210 million outpatient oral antibiotic prescriptions were given in the US annually (U.S. Centers for Disease Control and Prevention (CDC), 2021). Assuming even distribution, this is equivalent to over 60% of the U.S. population alone affected every year, contributing to the global rising trends of the aforementioned diseases.

Given its impact on host health, various approaches to replenish damaged gut microbiota have been attempted. Fecal microbiota transplant (FMT) is getting approval for treating *C. difficile* infections (U.S. Food and Drug Administration (FDA), 2019; Therapeutic Goods Administration, 2021), but the procedure was associated with acquired infections including a death by ESBL *E. coli* (FDA, 2019). It further led to a surge in resistome in both human and gnotobiotic pig recipients post FMT procedures (Liu and Wang, 2020). Autologous fecal microbiota transplantation (auto-FMT) facilitated the restoration of the gut microbiota and immune functions in patients (Schluter et al., 2020). Yet, reassessing the gut microbiota profiles and resistome illustrated that auto-FMT using own fecal microbiota banked before disruptive drug therapy, with the intention to reduce introduction of infectious agents from other donor(s), still resulted in the rise of resistome and opportunistic pathogens in multiple recipients even without further antibiotic treatment (Wang et al., unpublished data. Communicated with authors of MKSCC and agreed on the above conclusion in January 2021).

Fermented foods have emerged in recent years as a popular alternative to repair disrupted gut microbiota (Taylor et al., 2020; Wastyk et al., 2021; O’Connor, 2022). Wastyk et al. (2021) reported increased diversity of the gut microbiota of the human recipients by consuming diets high in fermented foods, but not by diets high in plant fibers. Fermented vegetables, followed by kombucha and fermented dairy products (yogurt and kefir) were among the most consumed categories of fermented foods by this group of subjects (Supplemental Data STable 1).

However, fermented foods rich in viable microbes are susceptible to AR. Before 2007, fermented dairy foods had been the most significant foodborne avenue transmitting AR to consumers (Wang et al., 2006). A gram of retail cheese contained up to 10^8^ copies of AR genes (Manuzon et al., 2007), and the *Bifidobacterium* strain supplemented to yogurt products worldwide contained a tetracycline (Tet) resistance-encoding gene (personal communication, 2007 ASM General Meeting). Various bacteria isolated from fermented foods with mobile AR genes were able to cause acquired resistance in human commensals and pathogens via horizontal gene transfer mechanisms (Wang et al., 2006; Li et al, 2011a; Jahan et al., 2016). Although successful mitigation of the AR gene pool was quickly achieved in mainstream fermented dairy products by 2011, primarily by removal of problematic fermentation starter cultures and probiotic strains from the product lines (Li et al, 2011b; Wang et al., 2019), other traditionally fermented products may still be prone to AR (Belletti et al., 2009; Comunian et al. 2010; Karasu et al., 2010; Nawaz et al., 2011; Pan et al., 2011; Zhou et al., 2012; Rebecchi et al., 2015; Zielińska et al., 2015; Nunes et al., 2016; Park et al., 2016; Guo et al. 2017; Erginkaya et al., 2018; Touret et al., 2018; Wang et al., 2018; Li et al., 2019; Leech et al., 2020; Yasir et al., 2022). Unlike mainstream modern dairy fermentation, which uses pasteurized milk and commercial starter cultures carefully screened by major culture companies, traditionally fermented vegetables and artisan cheeses, for instance, still rely on microbiota associated with raw materials, environment, or “mother” cultures other than carefully screened commercial starters (Belletti et al., 2009;

Comunian et al. 2010; Pan et al., 2011; Wang & Schaffner, 2011; Zielińska et al., 2015; Nunes et al., 2016; Park et al., 2016; Touret et al., 2018; Leech et al., 2020; Yasir et al., 2022).

The fermented food and beverage market is expected to grow by 5.6% in the next 10 years and will exceed US$ 989.2 billion by 2032 amid escalating demand for healthy and nutritious foods (Future Market Insights, 2022). The COVID-19 pandemic has further facilitated the explosive growth of sales of fermented products such as sauerkraut and kimchi (Manskar & Raskin, 2020). Given the booming market and increased consumption, re-assessing potential AR risks associated with traditionally fermented food products has become an urgent food safety and public health need, especially for susceptible populations with compromised digestive tract and immune systems. Therefore, this study investigated the incidences of antibiotic resistome in representative fermented foods and the impact of fermented food intervention on human gut resistome.

## Methods and Materials

### Host gut resistome assessment

In the original study by Wastyk et al. (2021), a total of 39 healthy adult participants were recruited, with 18 randomly assigned to the dietary intervention group by fermented foods and the remaining 21 assigned to the plant fiber intervention group. Three participants in the fiber intervention group dropped out of the study, thus both groups had 18 participants who finished the whole study (25 female,11 male, with an average age of 52 ± 11 years). Fecal shotgun metagenomics data of these participants were collected at four checkpoints, i.e., Week -2, Week 0, Week 8, and Week 10, throughout the study and deposited at the NCBI BioProject database (ID number: PRJNA743361). After initial screening for data availability, 3 participants in the fiber group and 1 participant in the fermented food group were dropped in our study due to missing Week 10 endpoint shotgun sequencing data in the database provided by the original research team. The raw sequencing data were retrieved from the database in Sequence Read Archive (.sra) format, converted to .fastq using the NCBI SRA Toolkit (version 3.0.0), and further processed on a high-performance supercomputer at the Ohio Supercomputer Center.

For quality control, FASTP tool (version 0.22.0) was used (parameters: -q 20 -u 20 -n 2 -l 80) to clean and trim the raw sequences before further annotation and analysis, as described by Chen et al. (2018). The cleaned sequences were then processed using the ARGs-OAP pipeline (Online Analysis Pipeline for Antibiotic Resistance Genes, version 2.5) (Yin et al., 2018). The resistome analysis criteria were set as hit length of 50%, e-value of 1e-07 and identity of 80%. The generated data were further analyzed and plotted using Tidyverse packages including ggplot2 (version 3.3.6) and dplyr (version 1.0.10), ggstatsplot package (version 0.10.0) and car package (Companion to Applied Regression, version 3.1-1) on R version 4.2.1 (Wickham et al., 2016; Fox & Weisberg, 2018; Wickham et al., 2019; Patil, 2021).

### Data transformation and analysis

The sums of the total ARG were calculated by adding up the relative abundances of all types of ARG annotated by ARGs-OAP. The percentage of total resistome change for each subject was calculated by the equation: (∑resistome reads after dietary intervention- ∑resistome reads before dietary intervention)/ ∑resistome reads before dietary intervention x 100%. The resistome data from Week -2 and Week 0 were averaged as the baseline to make sure all subjects have baseline data before dietary intervention and to reduce natural variation in gut microbiota. Normality and variance of data sets were verified using Shapiro-Wilk normality test and Levene’s test. Violin-boxplots with Student’s t-test p-values (α<0.05) were generated using ggstatsplot package (Patil, 2021). Other plots were constructed using the ggplot2 package (Wickham et al., 2016). Summary statistics such as mean, median and standard deviation and significance of dietary intervention were calculated using the dplyr package (Wickham et al., 2022).

### Source of fermented food samples

Representative kimchi samples of different brands were purchased from 7 retail stores including independent operations and national chains, 1 chain restaurant, and 3 local Japanese and Korean restaurants in Columbus, OH, from 2021–2023. The artisan cheese products were purchased from a national grocery chain store in Columbus, OH, with 4 purchased in 2021 and 4 in 2022. Sample designation and brief description were illustrated in Supplemental Data STable 2.

### Recovery and assessment of viable bacteria from fermented food samples

Five grams of each food sample were stomached in 45 ml 0.1% peptone water using a Seward stomacher 80 (Seward, UK). The juice was assessed for total bacteria and AR bacteria using Brain Heart Infusion (BHI), Luria-Bertani (LB) or De Man, Rogosa and Sharpe (MRS) agar plates with cycloheximide or nystatin as mold inhibitor, with or without the corresponding antibiotic (16 or 32 µg/mL ampicillin; 32 or 64 µg/mL tetracycline; 2 µg/mL erythromycin for BHI or MRS). A control sample with only peptone water was processed under the exact same condition and plated to make sure no contamination was introduced during the sample processing. The plates were incubated at 30°C, aerobically or anaerobically. Single colonies recovered from antibiotic-containing agar plates were picked based on representation in morphology and identified by Sanger sequencing of 16S rRNA gene PCR amplicons (Wang et al., 2006). The MIC of recovered AR isolates against 4 commonly used antibiotics were determined by a microdilution procedure (Stock et al., 2003), but using the corresponding recovery media instead. MIC of some representative AR isolates against a broader spectrum of antibiotics was further determined using commercial antimicrobial susceptibility kits Sensititre® (Thermo Scientific, USA, MA) GPN3F for Gram positive isolates and GN3F for Gram negative isolates, following the manufacturer’s instructions, with MRS or BHI broth replacing the Sensititre Mueller Hinton broth for better growth of each microbe. CLSI control strains *Escherichia coli* ATCC 25922 and *Enterococcus faecalis* 29212 (ATCC, Manassas, VA) were used as the MIC control standards.

### Antibiotic resistome of representative fermented food items

Twenty-five grams of food sample were mixed with 25 ml 0.1% peptone water and stomached as above. The large particles of food debris were removed with a sterile sieve. The liquid phase was centrifuged at 2500 relative centrifugal force by Multifuge X1R (Thermo Scientific, USA, MA) for 15 min. The pellet was washed once with 0.1% peptone water and subjected to total DNA extraction with a QIAamp PowerFecal Pro DNA Kit (Qiagen, Germany) following manufacturer’s instruction. Alternatively, the culture-recovered microbiota were scraped from agar plates with 3mL 0.1% peptone water and subjected to DNA extraction using the aforementioned QIAamp kit. The DNA samples were subjected to shotgun metagenomic sequencing (2X155bp) with average depth of 16.7 to 20 million paired-end reads for each sample, using an Illumina NextSeq2000 Sequencing System. The raw sequences were cleaned (parameters: -q 20 -u 40 -n 2 -l 80) and analyzed with the same metagenomic tools and parameters as described above to obtain relative abundance of AR genes normalized to the number of 16S rRNA gene. The sequencing data were further assessed for microbiota profiles through the Kraken2 pipeline with standard database and minimum-base-quality 20, as described previously (Wood et al., 2019). Food microbiota profile of Kimchi #7 was also assessed by 16S rRNA sequencing by an Illumina MiSeq system.

All local metagenomic data analyses were conducted on a high-performance supercomputer at the Ohio Supercomputer Center.

## Results

### Impact of food intervention on fecal microbiota resistome

Ten-week dietary interventions by diets high in fermented foods or plant fibers led to 11 increased and 6 decreased, as well as 10 increased and 5 decreased gut antibiotic resistome in human subjects, respectively, but the impact by the two types of diets was distinctive (Fig 1 & Fig 2). Fermented foods intervention led to significant changes in fecal antibiotic resistome of subjects with a p-value of 0.03, and the mean of resistome increased from 0.36 to 0.42 copies/16S rRNA gene (Fig 2A). Meanwhile, intervention by high plant fiber diets caused insignificant change in host gut resistome, with a p-value of 0.94. The mean of resistome remained as 0.38 copies/16S rRNA gene in subjects before and after dietary intervention (Fig 2B). The summarized resistome data used are illustrated in Supplemental Data STable 3. For details of the resistome data see supplemental data STable 9 & 10.

**Fig 1.**
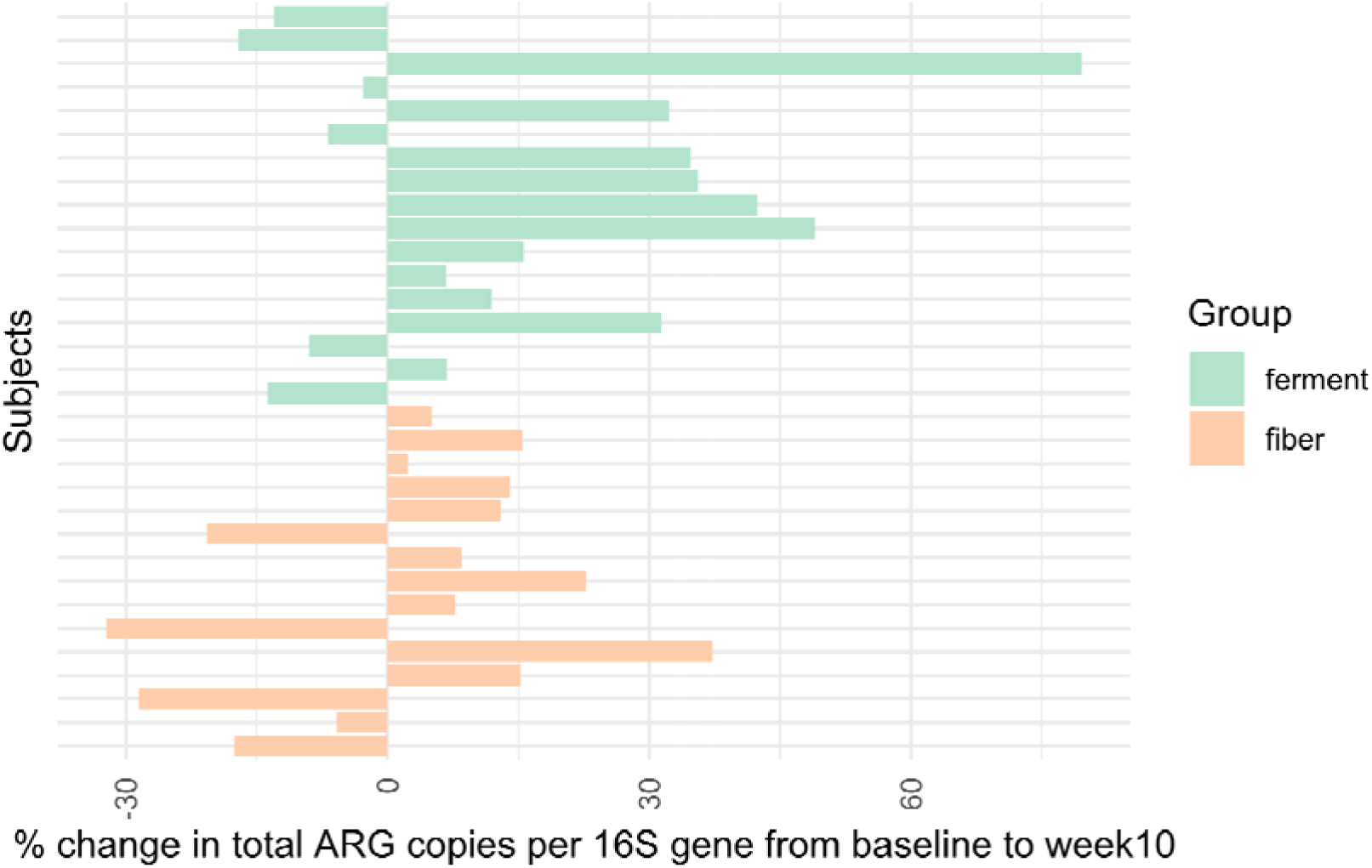
Impact of 10-week food intervention on the change of antibiotic resistome of fecal microbiota of human subjects in percentage comparing to the baseline. Green: by foods rich in fermented products; orange: by foods rich in plant fibers.

**Fig 2.**
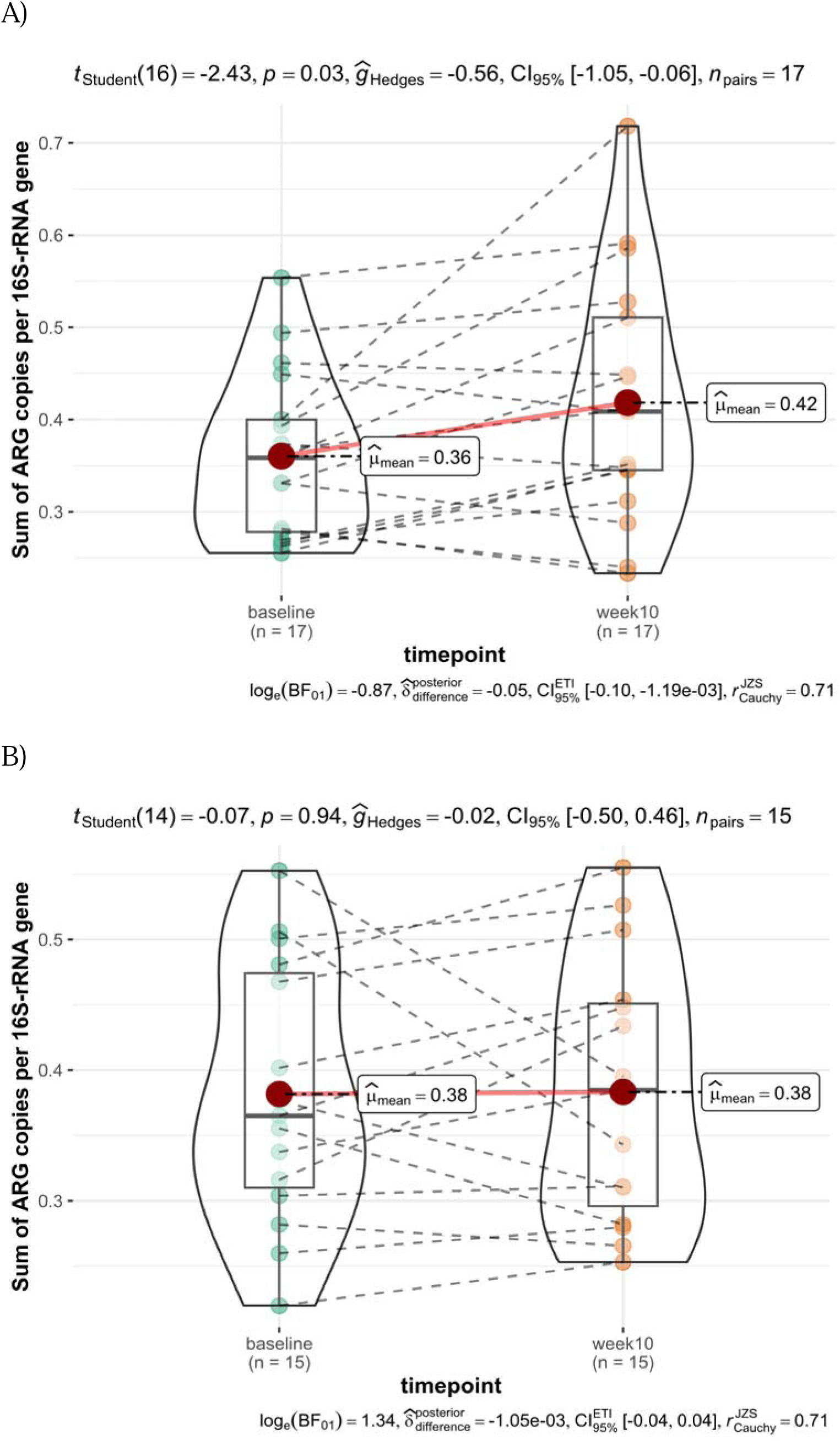
Violin-boxplots with Student’s t-test p-values illustrating the impact of 10-week food intervention on the change of antibiotic resistome of fecal microbiota of human subjects comparing to the baseline by diets rich in A) fermented foods; B) plant fibers.

### Antibiotic resistome of traditionally fermented foods

This pilot screening of retail fermented foods included various brands distributed nationwide through national, regional grocery chains and independent retailers. Multidrug resistance, bacitracin and macrolide-lincosamide-streptogramin (MLS) genes were most abundant in pooled retail kimchi samples (K1-K4) recovered from BHI plates (Fig 3A and Table 1), and they were also among the most abundant AR genes of all the 5 individual kimchi microbiota (K7-K11) assessed directly, despite the exact abundance of the top AR genes varied among these samples (Table 1 & Fig 4A). Likewise, even though AR isolates were identified in 9 out of 10 kimchi samples illustrated (K5, K7-K14, Table 2), the overall abundance of resistome of assessed samples also varied, ranging from 0.372 (K9) to 0.029 (K11) copies of AR genes/16S rRNA (Fig 4A).

**Fig 3.**
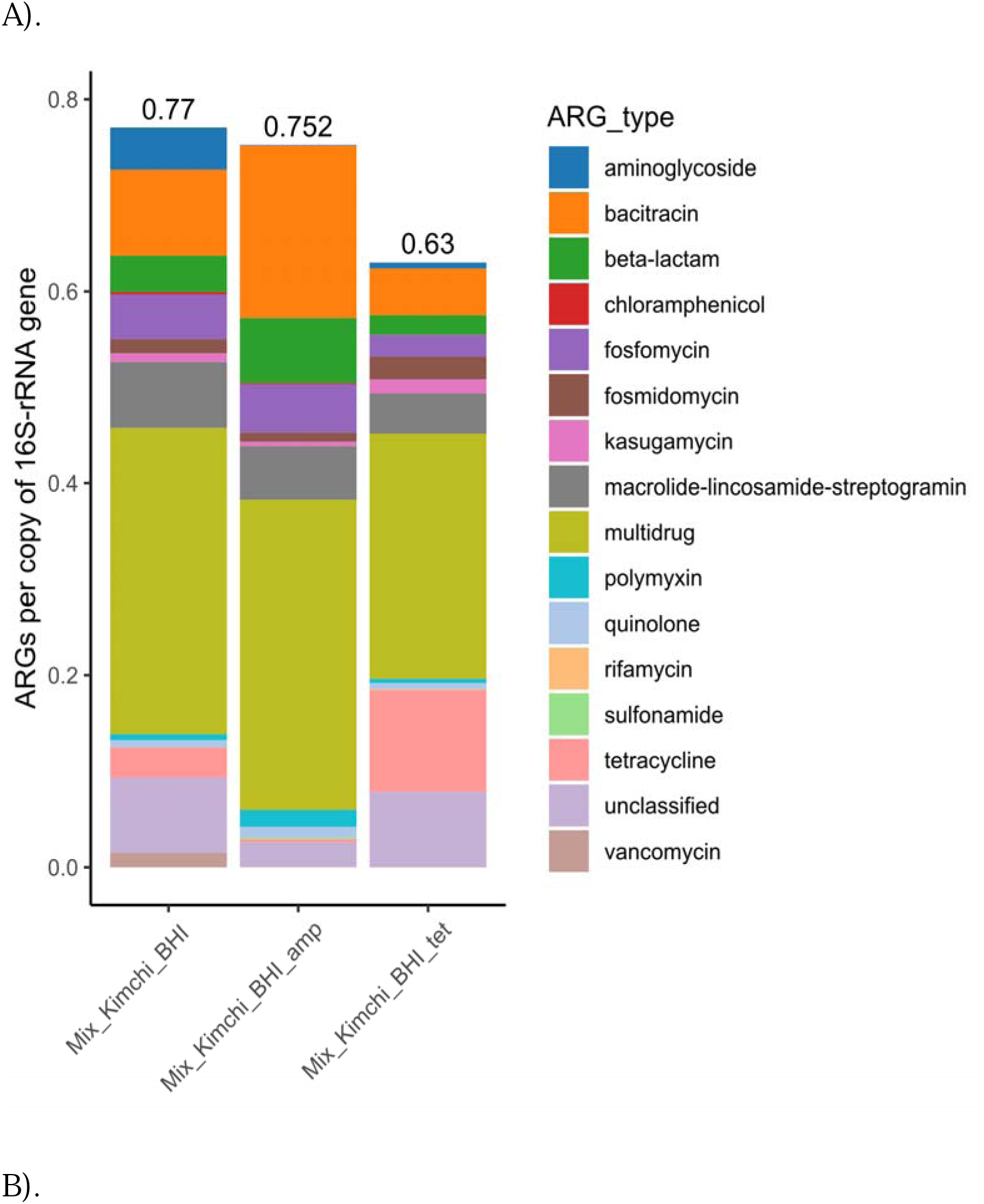
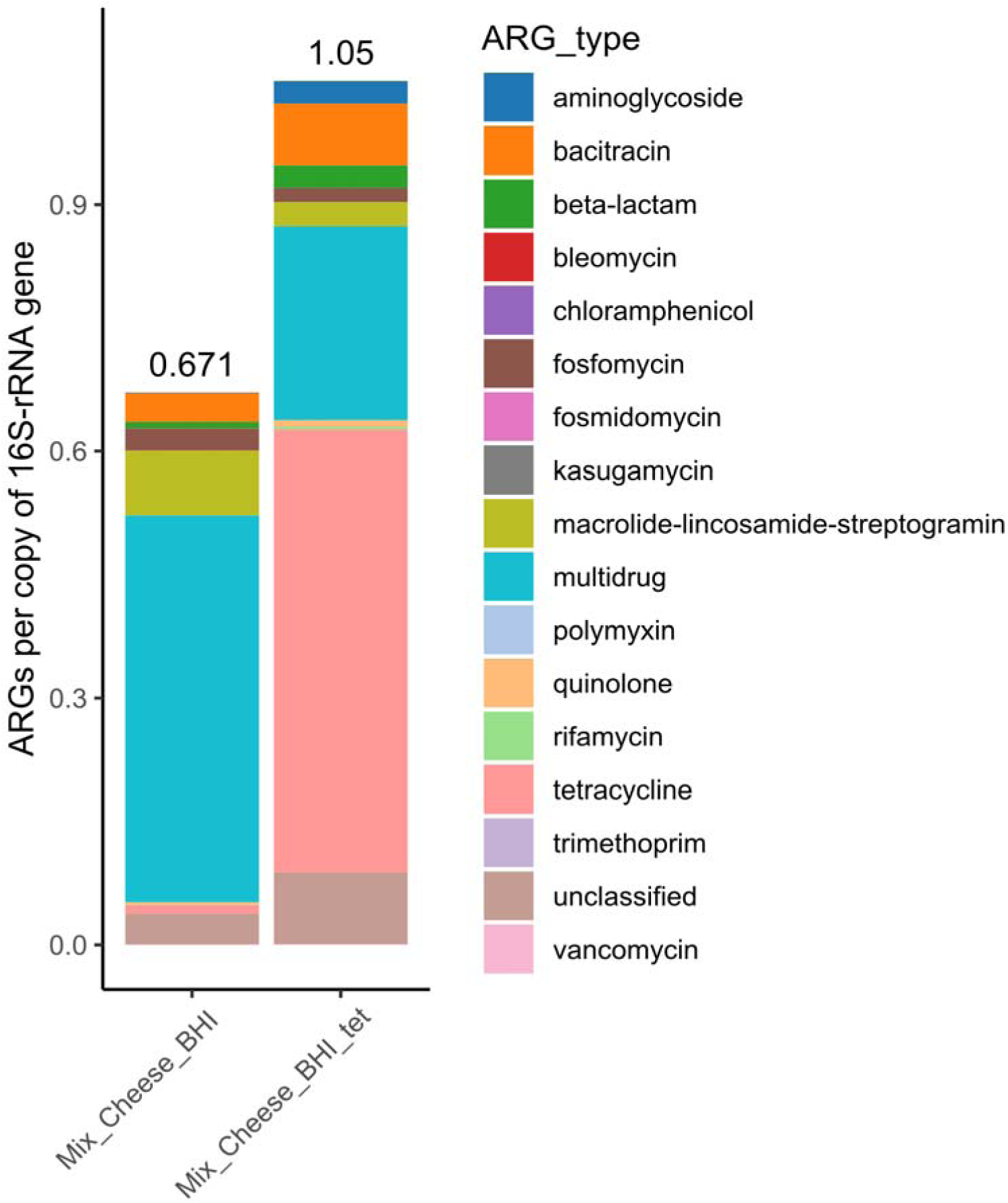
Antibiotic resistome of pooled traditionally fermented food microbiota recovered from agar plates. A) 4 kimchi samples (K1-K4) purchased in 2021. B) 4 artisan cheeses (C1-C4) purchased in 2021.

**Fig 4.**
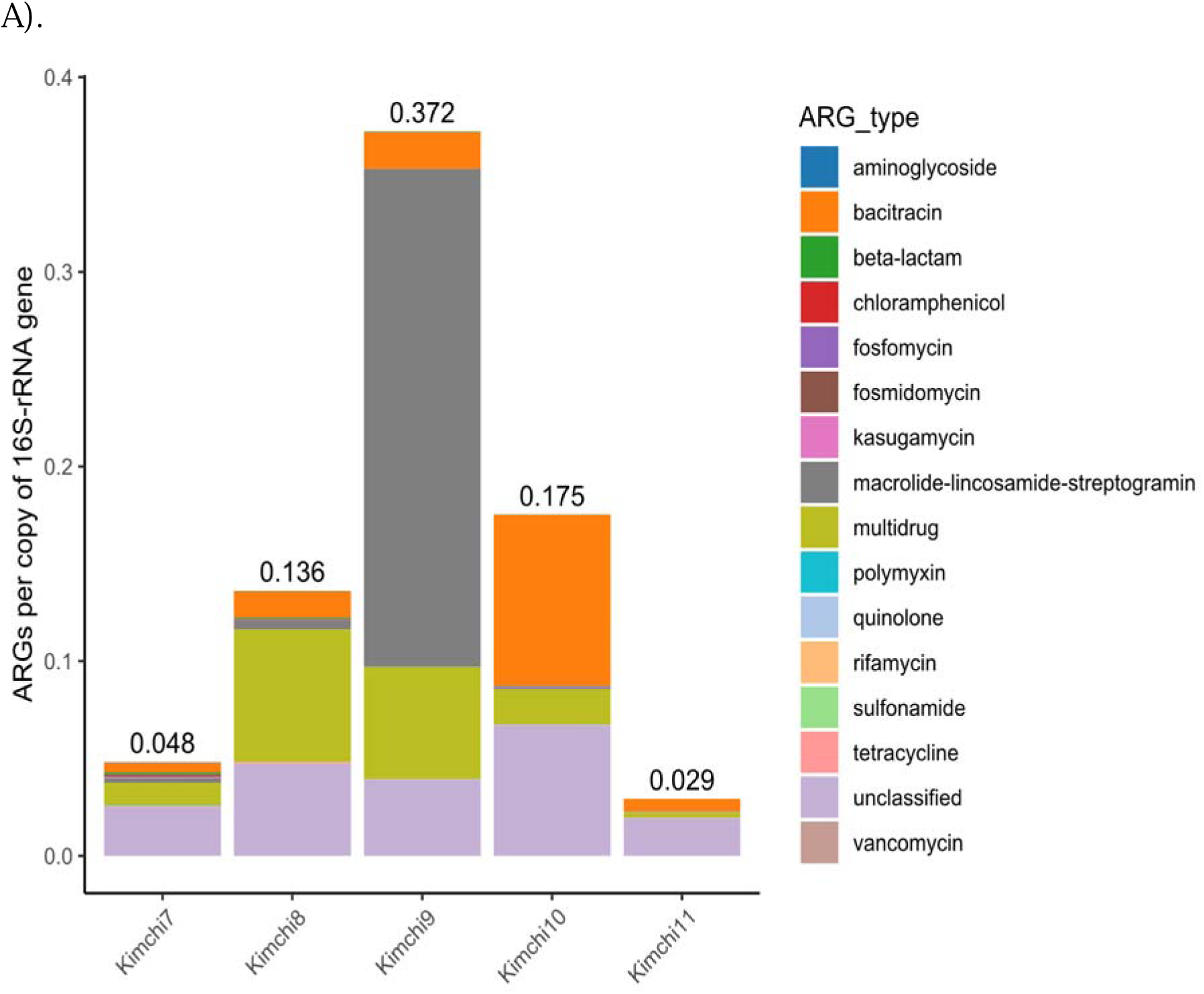
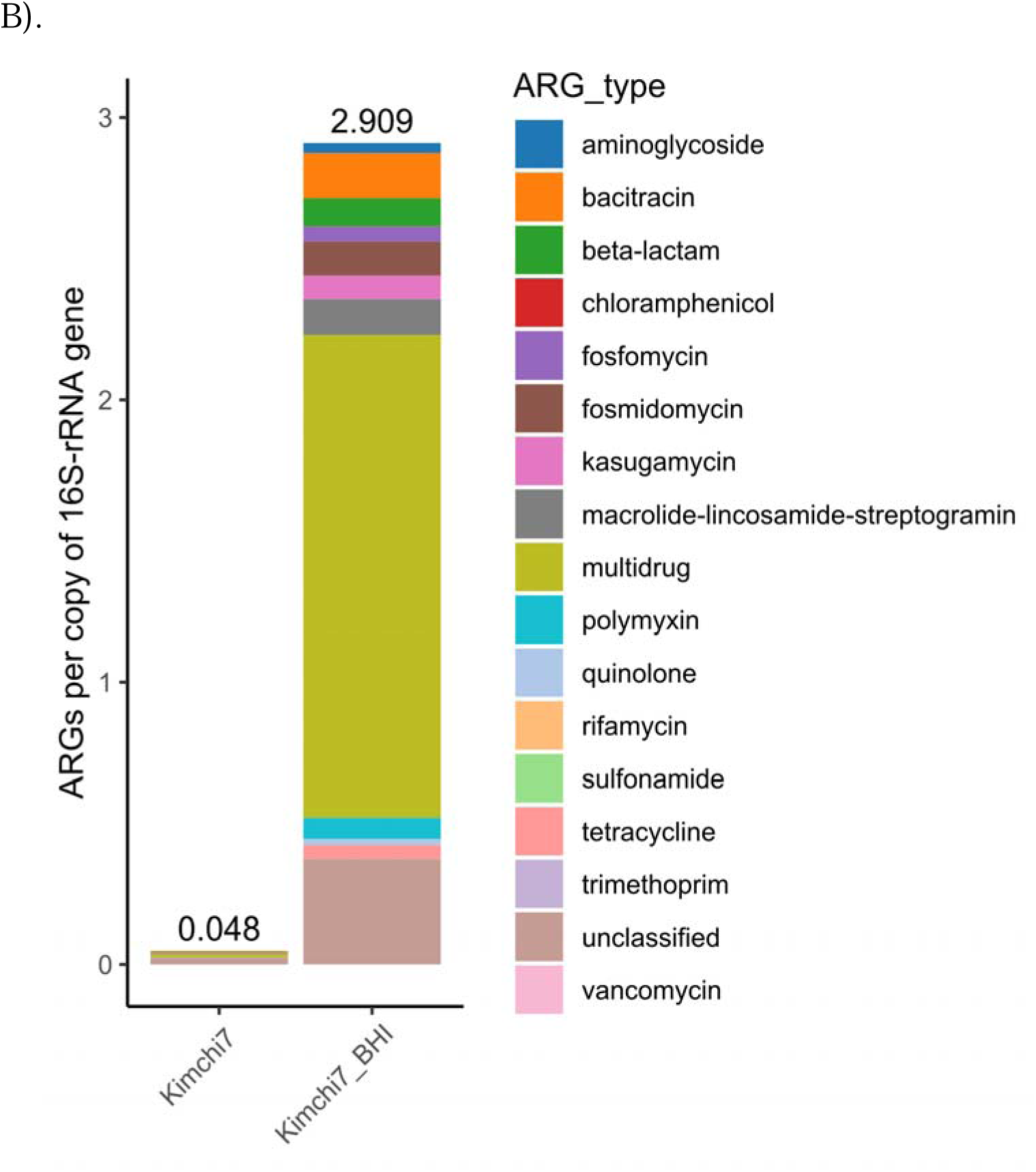
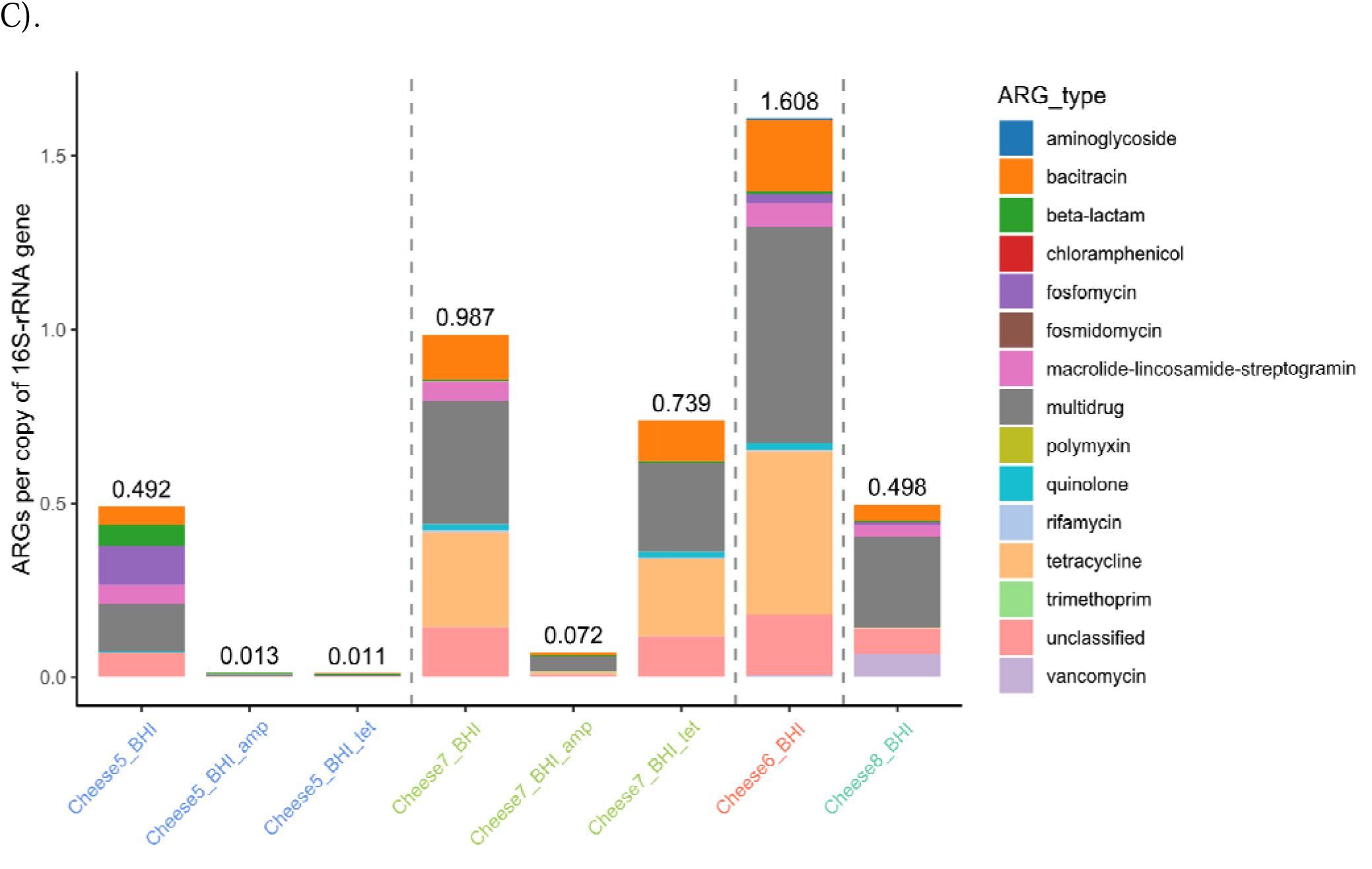
Resistome assessment of individual fermented foods purchased in 2022. A) direct resistome of kimchi samples; B) resistome of kimchi microbiota (left bar) and recovered BHI microbiota (right bar) of kimchi sample 7; C). resistome of BHI recovered cheese microbiota.

**Table 1.** Most abundant AR genes in kimchi samples*.

| Sample # | Rank |  |  |  |  |
| --- | --- | --- | --- | --- | --- |
|  | 1 | 2 | 3 | 4 | 5 |
| K1-4 (BHI, pooled) | Multidrug<br>0.3189* | Bacitracin<br>0.0901 | Unclassified<br>0.0780 | MLS**<br>0.0684 | Fosfomycin<br>0.0465 |
| K1-4 (BHI-Tet, pooled) | Multidrug<br>0.2556 | Tetracycline<br>0.1059 | Unclassified<br>0.0777 | Bacitracin<br>0.0485 | MLS<br>0.0422 |
| K1-4 (BHI-Amp, pooled) | Multidrug<br>0.3235 | Bacitracin<br>0.1795 | β-lactam<br>0.0680 | MLS<br>0.0557 | Fosfomycin<br>0.0498 |
| K7 (BHI) | Multidrug<br>1.7085 | Unclassified<br>0.3728 | Bacitracin<br>0.1609 | MLS<br>0.1278 | Fosmidomycin<br>0.1216 |
| K7 | Unclassified<br>0.0248 | Multidrug<br>0.0118 | Bacitracin<br>0.0050 | MLS<br>0.0023 | Fosmidomycin<br>0.0012 |
| K8 | Multidrug<br>0.0681 | Unclassified<br>0.0469 | Bacitracin<br>0.0136 | MLS<br>0.0047 | Tetracycline<br>0.0006 |
| K9 | MLS<br>0.2557 | Multidrug<br>0.0577 | Unclassified<br>0.0390 | Bacitracin<br>0.0191 | Vancomycin<br>0.0001 |
| K10 | Bacitracin<br>0.0880 | Unclassified<br>0.0669 | Multidrug<br>0.0182 | MLS<br>0.0009 | Fosfomycin<br>0.0005 |
| K11 | Unclassified<br>0.0191 | Bacitracin<br>0.0063 | Multidrug<br>0.0026 | Tetracycline<br>0.0002 | MLS*<br>0.0002 |

\*Numerical readings of AR: in total ARG copies per 16S gene.
\*\*MLS: Macrolide-Lincosamide-Streptogramin.

**Table 2.**
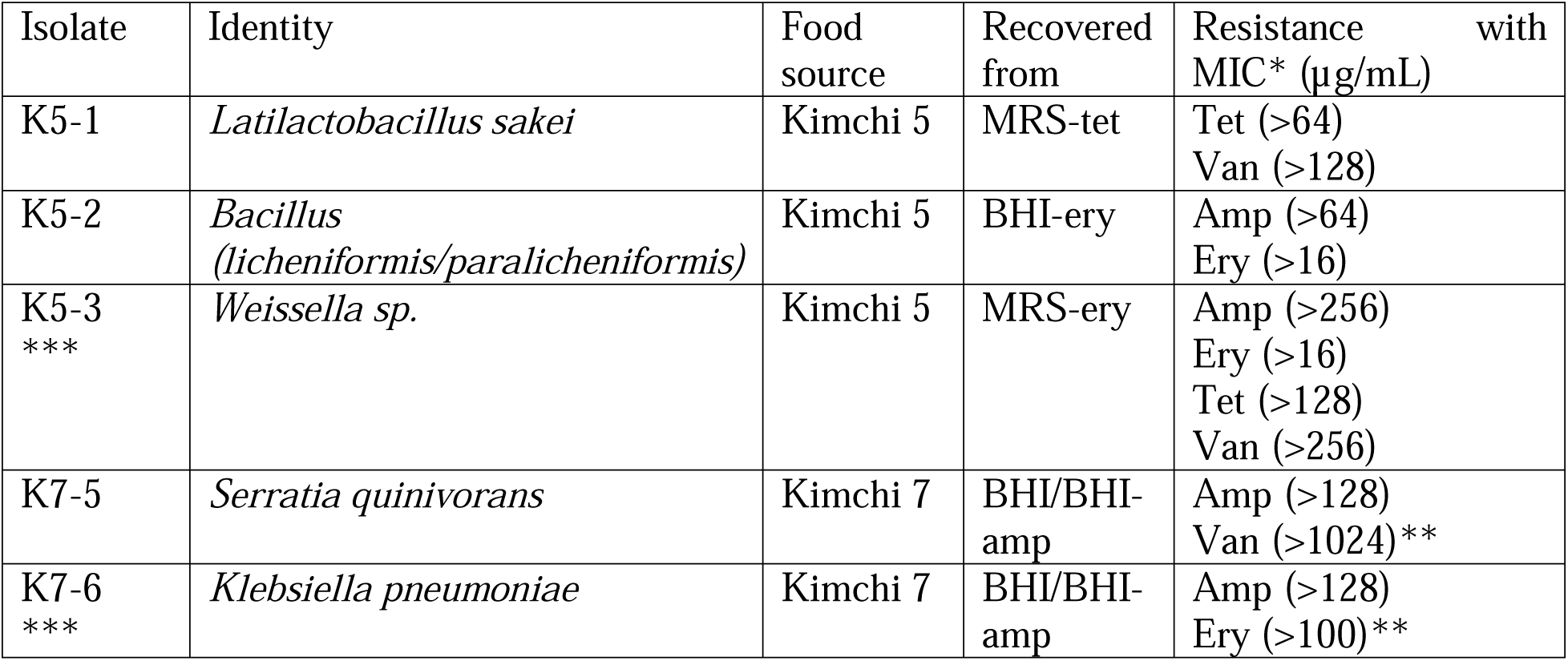
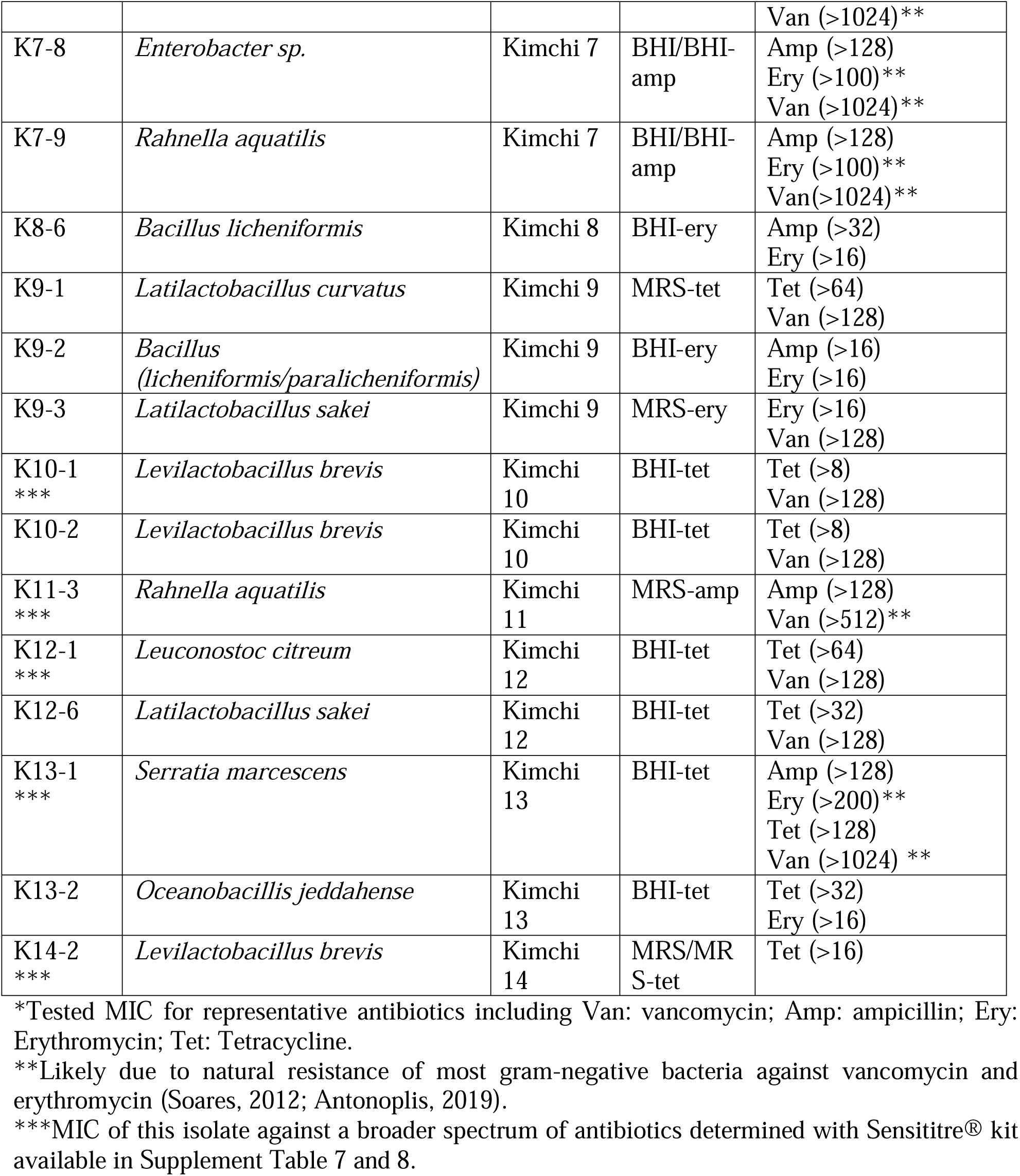
Summary of identified AR colonies from kimchi products purchased in 2022 and 2023*.

Figure 4B further compares the resistome outcomes of kimchi sample K7 using total DNA extracted from kimchi microbiota directly (0.048 copies of AR genes/16S ) and total DNA of kimchi microbiota recovered from BHI agar plate (2.909 copies of AR genes/16S). The results indicated that with specific medium and incubation condition, even without antibiotic selective pressure, certain bacteria of the kimchi microbiota might have been selectively enriched, leading to the drastic difference in detected antibiotic resistome. Likewise, the top genera of pooled kimchi microbiota recovered from BHI plates (Supplemental Data STable 4) were affected by cultivation conditions too, with and without antibiotics, as the illustrated resistome (Fig 4B).

Indeed, Table 3 illustrates that the top genera of each individual kimchi microbiota (K7 to K11) assessed using total DNA extracted directly were dominated by lactic acid bacteria well-recognized for driving kimchi fermentation. The data further demonstrated that antibiotic resistome and AR bacteria were still prevalent in these successfully fermented kimchi samples.

**Table 3.** Top 5 genera classified of microbiota directly of 5 kimchi samples purchased in 2022.

| Sample | Top genera | Percentage |
| --- | --- | --- |
| Kimchi7 | *Leuconostoc* | 94.87% |
|  | Other | 3.17% |
|  | *Rahnella* | 0.85% |
|  | *Weissella* | 0.48% |
|  | *Levilactobacillus* | 0.36% |
|  | *Xanthomonas* | 0.27% |
| Kimchi8 | *Latilactobacillus* | 55.98% |
|  | *Leuconostoc* | 28.90% |
|  | *Lactiplantibacillus* | 7.49% |
|  | Other | 5.46% |
|  | *Lactococcus* | 1.59% |
|  | *Levilactobacillus* | 0.58% |
| Kimchi9 | *Latilactobacillus* | 81.60% |
|  | *Leuconostoc* | 11.79% |
|  | Other | 3.45% |
|  | *Weissella* | 2.02% |
|  | *Lactiplantibacillus* | 0.59% |
|  | *Levilactobacillus* | 0.55% |
| Kimchi10 | *Lactiplantibacillus* | 54.60% |
|  | *Leuconostoc* | 32.89% |
|  | *Levilactobacillus* | 9.80% |
|  | Other | 1.86% |
|  | *Weissella* | 0.55% |
|  | *Latilactobacillus* | 0.30% |
| Kimchi11 | *Weissella* | 58.85% |
|  | *Leuconostoc* | 39.19% |
|  | Other | 1.41% |
|  | *Latilactobacillus* | 0.23% |
|  | *Lactiplantibacillus* | 0.19% |
|  | *Stenotrophomonas* | 0.13% |

The resistome results of pooled artisan cheese samples C1 to C4 (Fig 3B) and individual cheese samples of C5 to C8 (Fig 4C), all using microbiota recovered from BHI agar plates, further illustrated the prevalence of various AR genes in these products. Multidrug, MLS and bacitracin were among the top AR genes of the pooled samples C1-C4, while multidrug, bacitracin, tetracycline and MLS were most abundant among individual samples C5 to C8 (Table 4). It is worth noting that the presented abundance of resistome of cheese microbiota recovered from BHI plates may also be affected by the medium and culture conditions, deviated from those of the original samples. Supplemental Data STable 6 further illustrated that *Staphylococcus* (including *Mammaliicoccus,* also known as *Staphylococcus*) was the top dominant genus of individual artisan cheese microbiota recovered from BHI agar plates.

**Table 4.** Most abundant AR genes in BHI plates recovered microbiota of artisan cheese samples.

| Sample # | Rank |  |  |  |  |
| --- | --- | --- | --- | --- | --- |
|  | 1 | 2 | 3 | 4 | 5 |
| C1-4 (BHI, pooled) | Multidrug<br>0.4704* | MLS**<br>0.0783 | Bacitracin<br>0.0353 | Fosfomycin<br>0.0265 | Tetracycline<br>0.0105 |
| C1-4 (BHI-Tet, pooled) | Tetracycline<br>0.5377 | Multidrug<br>0.2359 | Unclassified<br>0.0876 | Bacitracin<br>0.0744 | MLS<br>0.0295 |
| C1-4 (BHI-Amp, pooled) | Multidrug<br>0.0308 | Tetracycline<br>0.0093 | MLS<br>0.0084 | Bacitracin<br>0.0053 | Unclassified<br>0.0033 |
| C5 (BHI) | Multidrug<br>0.1343 | Fosfomycin<br>0.1123 | Unclassified<br>0.0709 | β-lactam<br>0.0617 | MLS<br>0.0555 |
| C5 (BHI-Tet) | Multidrug<br>0.0033 | β-lactam<br>0.0031 | Tetracycline<br>0.0014 | Bacitracin<br>0.0012 | Unclassified<br>0.0012 |
| C5 (BHI-Amp) | Multidrug<br>0.0040 | β-lactam<br>0.0038 | Unclassified<br>0.0015 | Bacitracin<br>0.0014 | MLS<br>0.0013 |
| C6 (BHI) | Multidrug<br>0.6202 | Tetracycline<br>0.4674 | Bacitracin<br>0.2046 | Unclassified<br>0.1750 | MLS<br>0.0694 |
| C7 (BHI) | Multidrug<br>0.3538 | Tetracycline<br>0.2740 | Unclassified<br>0.1442 | Bacitracin<br>0.1308 | MLS<br>0.0557 |
| C7 (BHI-Tet) | Multidrug<br>0.2558 | Tetracycline<br>0.2215 | Unclassified<br>0.1183 | Bacitracin<br>0.1176 | Quinolone<br>0.0167 |
| C7 (BHI-Amp) | Multidrug<br>0.0394 | Bacitracin<br>0.0085 | Tetracycline<br>0.0078 | Unclassified<br>0.0075 | β-lactam<br>0.0035 |
| C8 (BHI) | Multidrug<br>0.2605 | Unclassified<br>0.0737 | Vancomycin<br>0.0671 | Bacitracin<br>0.0472 | MLS<br>0.0345 |

\*Numerical readings in total ARG copies per 16S gene.
\*\*MLS: Macrolide-Lincosamide-Streptogramin.

### Representative AR bacteria from traditionally fermented foods

Various AR bacteria were recovered from traditionally fermented products. Under very limited cultivation conditions applied in this study, AR bacteria were identified from 9 out of 10 kimchi and 4 out of 4 cheese samples assessed by culture recovery (Table 2 & 5).

Identified AR isolates from kimchi products ranged from important human pathogens to organisms still considered as commensals, as well as lactic acid bacteria driving food fermentation. For instance, AR opportunistic pathogens such as *Klebsiella pneumoniae* and *Serratia marcescens* are recognized agents for nonsocomial infections (Farmer, 2003; Mahlen, 2011). *Rahnella aquatilis* as a human pathogen can cause bacteremia, sepsis, urinary tract infection, etc. As illustrated in Table 2 and Supplementary Table 7, kimchi isolates *Klebsiella pneumoniae* K7-6 and *Rahnella aquatilis* K11-3 are resistant to multiple antibiotics. Particularly, *Serratia marcescens* K13-1 is highly resistant to almost all key antibiotics (21 out of 22), including the 4^th^ generation of cephalosporin antibiotic as well as carbapenem antibiotics, with MICs exceeded the highest detection limit of 12 antibiotics by the Sensititre assessment (Supplemental Table 7). In addition, isolates of several species of *Latilactobacillus*, *Leuconostoc* and *Weissella*, commonly involved in driving kimchi fermentation, were among those highly resistant to tetracycline, erythromycin, ampicillin and vancomycin, and some isolates exhibited multidrug resistance (Table 2). *Weissella* K5-3 further exceeded the highest MIC levels of all 18 antibiotics examined using the Sensititre panel (Supplemental Table 8).

Identified AR isolates from cheese products were mostly *Staphylococcus* spp. resistant to tetracycline or erythromycin, with one isolate also resistant to vancomycin. An isolate of *Enterococcus* sp. was highly resistant to vancomycin and could be classified as vancomycin resistant *Enterococcus* (Rengaraj, 2016). It is also multidrug-resistant (Supplemental Table 8).

### Antibiotic exposure further shaped the profiles of fermented food originated microbiota

As illustrated in Fig. 5 and Supplemental Table 6, cheese microbiota harvested from BHI plates without antibiotics were dominated by *Mammaliicoccus* spp. (formerly *Staphylococcus*), *Staphyloccocus, Camobacterium*, *Psychrobacter* and *Glutamicibacter (*belonging to *Micrococcaceae*) for cheese sample C5. But at the presence of Tet or Amp, the dominant bacteria switched to *Stenotrophomonas*, *Pseudomonas*, *Burkholderia*, *Alcanivorax* and *Streptococcus*. Likewise, while the dominant cultures for cheese C7 recovered on BHI were *Staphylococcus* and *Lactococcus*, the top ranked bacteria of cheese originated microbiota shifted to *Pseudomonas, Burkholderia*, *Staphylococcus, Streptococcus* and *Leuconostoc* on BHI-Amp, and *Staphylococcus* on BHI-Tet. In the case of C7 and pooled sample C1-C4, the cheese starter culture *Lactococcus* was essentially eliminated when the cheese microbiota was exposed to either of the two antibiotics.

**Fig 5.**
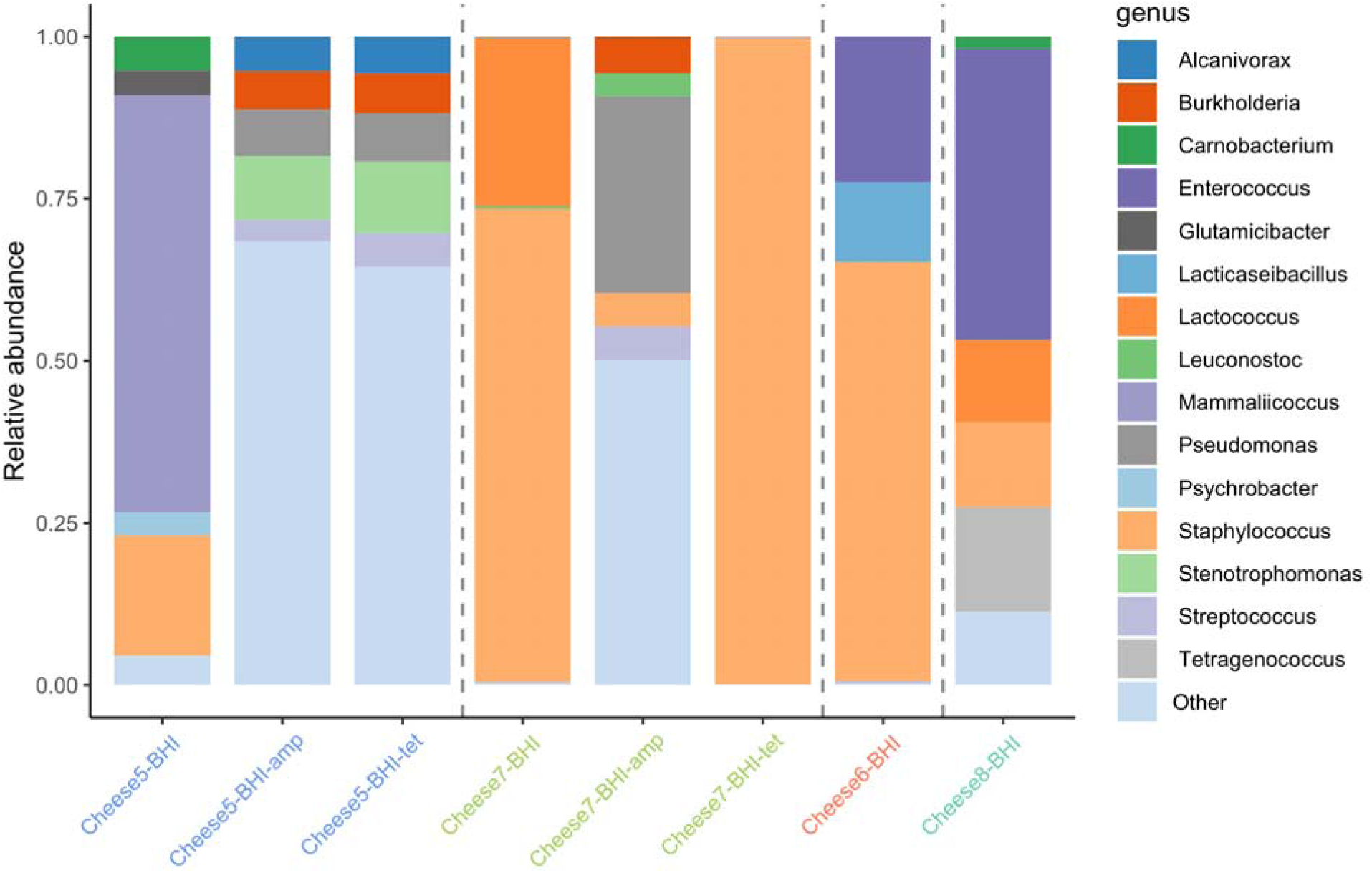
Detected cheese microbiota recovered from BHI agar plates with and without antibiotics.

Although antibiotic-containing agar plates were used to screen for AR bacteria, the above result also indicated how the food-originated microbiota might impact gut microbiota when the hosts received antibiotic treatment. *Stenotrophomonas*, *Pseudomonas*, *Burkholderia,* and *Staphylococcus* are recognized opportunistic pathogens associated with antibiotic resistant hospital acquired infections, and most *Streptococcus* species are pathogens.

Microbial profiling by metagenomics of dominant cheese bacteria recovered on BHI plate included *Staphylococcus, Enterococcus* and *Lactocaseibacillus* (formerly *Lactobacillus*) for cheese sample C6, as well as *Enterococcus, Tetragenococcus, Staphylococcus, Lactococcus* and *Camobacterium* for sample C8, respectively (Fig. 5 and Supplemental Table 6). In agreement, some of the mentioned antibiotic resistant bacteria such as multidrug resistant *Enterococcus* sp. and *Staphylococcus* were also confirmed in these samples (Table 5).

**Table 5.**
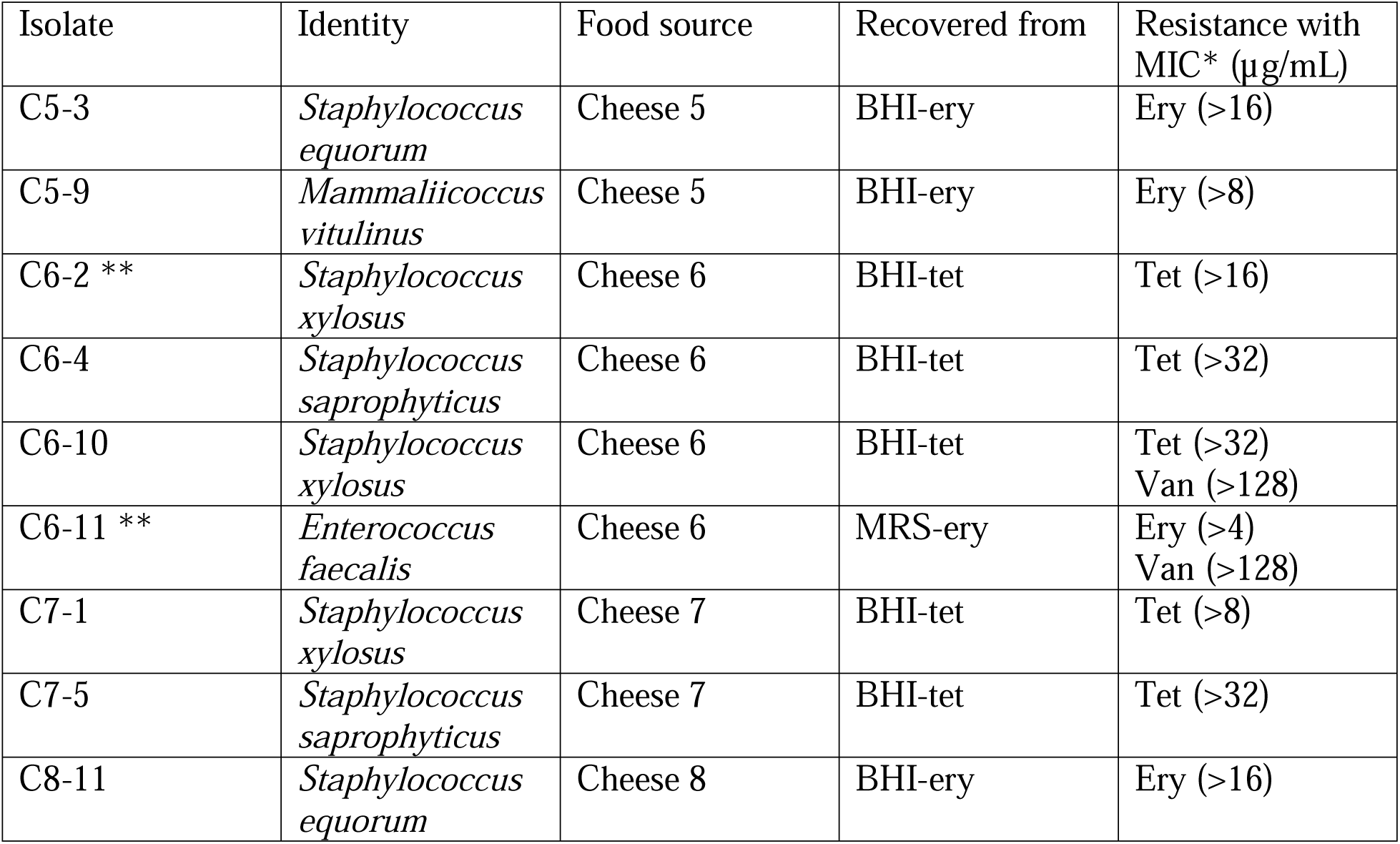
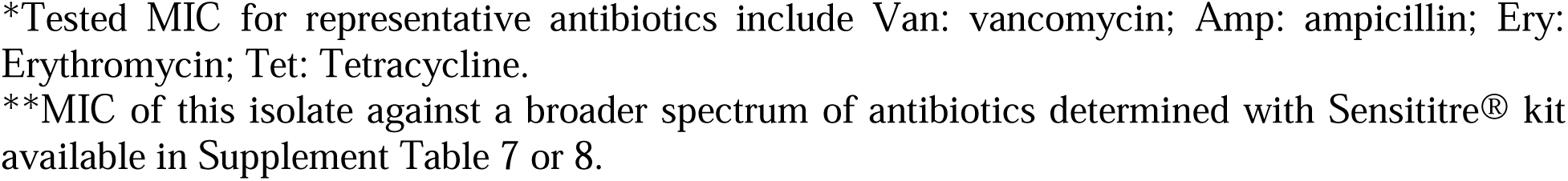
Summary of identified AR colonies from artisan cheese products purchased in 2022*.

| Isolate | Identity | Food source | Recovered from | Resistance with MIC* (μg/mL) |
| --- | --- | --- | --- | --- |
| C5-3 | *Staphylococcus equorum* | Cheese 5 | BHI-ery | Ery (>16) |
| C5-9 | *Mammaliococcus vitulinus* | Cheese 5 | BHI-ery | Ery (>8) |
| C6-2 ** | *Staphylococcus xylosus* | Cheese 6 | BHI-tet | Tet (>16) |
| C6-4 | *Staphylococcus saprophyticus* | Cheese 6 | BHI-tet | Tet (>32) |
| C6-10 | *Staphylococcus xylosus* | Cheese 6 | BHI-tet | Tet (>32)<br>Van (>128) |
| C6-11 ** | *Enterococcus faecalis* | Cheese 6 | MRS-ery | Ery (>4)<br>Van (>128) |
| C7-1 | *Staphylococcus xylosus* | Cheese 7 | BHI-tet | Tet (>8) |
| C7-5 | *Staphylococcus saprophyticus* | Cheese 7 | BHI-tet | Tet (>32) |
| C8-11 | *Staphylococcus equorum* | Cheese 8 | BHI-ery | Ery (>16) |

\*Tested MIC for representative antibiotics include Van: vancomycin; Amp: ampicillin; Ery: Erythromycin; Tet: Tetracycline.
\*\*MIC of this isolate against a broader spectrum of antibiotics determined with Sensititre® kit available in Supplement Table 7 or 8.

## Discussion and Conclusion

Although unexpected for many, the susceptibility of fermented foods to AR has been well-recognized at least in the food microbiology community. Since the first systematic demonstration of the problem in the early 2000s with a broad spectrum of AR isolates, including starter cultures, opportunistic pathogens and commensals of mainstream fermented dairy products being identified and characterized (Wang et al., 2006), AR bacteria have further been isolated from various fermented foods worldwide (Muñoz et al., 2014; Fraqueza, 2015; Kim et al., 2021). For instance, *Pantoea agglomerans* (formerly *Enterobacter agglomerans*, or *Erwinia herbicola*), an opportunistic pathogen causative to a wide range of opportunistic infections, especially in immunocompromised patients, was isolated from kimchi in South Korea. The genome of *P. agglomerans* isolate KM1 contained 13 genes conferring resistance to clinically important antibiotics, and the strain exhibited immunostimulatory properties *in vitro*, including the production of pro-inflammatory and anti-inflammatory cytokines in stimulated cells (Guevarra et al., 2021).

This study, however, using a combination of approaches, illustrated the high prevalence and the abundance of antibiotic resistome in popular retail traditionally fermented foods. Although the retail products were purchased in Columbus Ohio, the products were mostly made in the US and distributed nationwide through independent stores and grocery chains. Therefore, they serve as a good indication for the prevalence of AR in similar products nationwide. Our team has further conducted traditional kimchi fermentation carefully in microbial controlled setting and has concluded that AR is inevitable in the final products due to the AR bacteria associated with the raw vegetable materials. This is consistent with the principle of fermentation that the natural microbiota from raw materials drive microbial succession in natural fermentation.

The original human study on dietary intervention by Wastyk et al. (2021) had a total of 36 healthy adult subjects completed the study, half received fermented foods and the other half dietary fiber intervention. Among those, the fecal microbiome outcomes of 32 subjects with required data points, including 17 subjects with fermented foods intervention and 15 subjects with diet high in plant fiber for comparison, were assessed and reported for the impact of the dietary intervention on host gut resistome in this study.

The sample size was still small, giving the diversities in human subjects and activities. Nevertheless, the illustration of antibiotic resistome in kimchi and artisan cheese microbiome, along with the high prevalence of confirmed AR bacteria in kimchi and artisan cheeses assessed in this pilot study, support the conclusion that consuming these traditionally fermented products would result in the rise of gut antibiotic resistome in consumers. Since only limited conditions were applied in cultivating AR bacteria in this pilot study, the recovered isolates in fact only represent a small percentage of the AR microbiota of the fermented foods. The pivotal impact of foodborne AR on host gut antibiotic resistome has previously been illustrated using the mice model by Zhang et al. (2013), as oral antibiotic treatment only led to the surge of the targeted AR gene pool in host gut microbiota when the mice had prior oral seeding of AR bacteria. Results from this pilot study thus call for more extensive investigations on the food safety risks associated with traditionally fermented foods, and for effective mitigation strategies to protect public health.

It is further worth noting that even in subjects who consumed only yogurt and kombucha (Subject 8020) or yogurt, kombucha and kefir (Subject 8014) (STable 3), the gut resistome of the subjects still increased 11% and 17%, respectively, after the 10-week intervention. The data suggests that consuming these products at least did not reduce the gut antibiotic resistome in the subjects.

The identification of AR bacteria from traditionally fermented products, ranging from human and plant pathogens to lactic acid bacteria driving food fermentation, especially those exhibited multidrug resistance and with extremely high MICs against antibiotics of clinical significance, is particularly concerning. Multidrug resistant *Klebsiella pneumoniae, Serratia marcescens,* as well as *Stenotrophomonas*, *Pseudomonas*, *Burkholderia, Staphylococcus* and *Streptococcus* are also known for causing hospital-acquired infections, including some of the most troublesome cases in healthcare facilities. Data from this study thus also provide an alternative interpretation on the route of dissemination and potential origins of infections in patients. For instance, *Serratia marcescens* K13-1 is extremely resistant to most antibiotics assessed by the Sensititre panel for Gram-negative bacteria, including the carbapenem antibiotics Meropenem and Ertapenem that are usually reserved for treating multidrug resistant bacterial infections (Supplemental Table 7). *Weissella s*pp. K5-3 is further highly resistant to all 18 antibiotics assessed by the Sensititre panel for Gram positive bacteria including the 3^rd^ generation of cephalosporin antibiotic Ceftriaxone (Supplemental Table 8). As lactic acid bacteria, *Weissella* spp. are known for driving vegetable fermentation and thus likely survive well even in successfully fermented products. Revealing the genetic elements responsible for the unusual AR profiles and their potential involvement in horizontal gene transfer will provide further insights on the food safety and public health risks associated with these AR bacteria.

Balanced gut microbiota as well as effective mucosal, intestinal epithelial and gut vascular barriers as part of an integral and functional gastrointestinal tract system are essential to host health (Gao et al., 2022). Physical damages to the intestinal barriers further have various health consequences. Intestinal tract problems such as inflammatory bowel diseases, enteric infections or overgrowth by pathogens such as EHEC and Shiga toxin-producing *E. coli*, *Shigella*, *Salmonella*, *Campylobacter*, *C. difficile*, enteric viruses, certain drug treatments such as the antibiotic clindamycin etc., even alcohol consumption can all cause acute or chronic intestinal barrier damages. Once the intestinal barriers collapse, intestinal microorganisms and their metabolites can get into the bloodstream, reach other organs and cause further health damages. Once entering the bloodstream, even commensals and probiotics can be problematic. For instance, the most popular probiotic strains *Bifidobacterium* and *L. rhamnosus* GG have still been found causative to bacteremia and sepsis in high-risk populations, with strain confirmation by metagenomic sequencing (Land et al., 2005; Boyle et al., 2006; Ohishi et al., 2010; Kochan et al., 2011; Bertelli et al., 2015; Angurana et al., 2018; Chiang et al., 2021; Aydoğan et al., 2022; Kulkarni et al., 2022; Colautti et al., 2022). While bacteremia and sepsis by commercial probiotic strains absence of AR genes are still controllable by mainstream antibiotics such as ampicillin, introducing the broad spectrum of AR bacteria highly resistant to clinically important antibiotics associated with traditionally fermented foods presents an underestimated and potentially serious public health risk, especially to targeted populations suffering from various gut symptoms associated with “leaky gut” as well as compromised immune functions. Bacteremia and sepsis risk, antibiotic therapy failures, and further gut microbiota destruction post-antibiotic treatment may significantly increase in the targeted susceptible consumer populations who intend to repair damaged gut microbiome by enhancing fermented food consumption.

With the recipients of over 210 million oral antibiotic prescriptions added to the susceptible population suffering from broad gut microbiota destruction annually in the US alone (CDC, 2021), the impacted patient population worldwide is astonishing. While various gut microbiota-replenishing approaches including fermented food intervention have been practiced desperately, they all have introduced additional public health risks to the patients. Results from this study thus further call for strategic efforts to tackle the key and shared driver of the antibiotic resistome surge and gut microbiota disruption in human and animals, by mitigating the applications of gut-impacting antibiotics (oral administration and biliary excretion) and offering alternatives, to significantly minimize unnecessary damages to gut microbiome for productive outcomes.

## Supporting information

Supplemental Table 9 resistome

Supplemental Table 10 resistome

## Acknowledgement

The study was possible with funding from the OSU Women and Philanthropy. YL and SF were funded by OSU FST Graduate Research Associateship for HW. YL is also the recipient of the 2022 Elwood Caldwell graduate fellowship of the Institute of Food Technologists (IFT). We thank Dr. Erin DiCaprio of UC Davis for sharing information on the retail trends in fermented foods. We also thank our collaborators who supported and enabled this research.

We also thank Dr. Justin Sonnenburg’s team for properly organizing and depositing the human study data published in Cell (2021) in the public domain, which enabled our data mining and reassessment.

Part of the data was presented at the 2022 ASM Microbe Satellite/Sino-Micro annual meeting (Washington, DC) and USDA multistate NC1206 project meeting on antibiotic resistance (Ames, IA).

## Data availability statement

All metagenomic sequencing data were deposited at the NCBI BioProject database (ID number: PRJNA944373).

