## Supplemental Table 1 to 6 for "High prevalence of antibiotic resistance in traditionally fermented foods as a critical risk factor for host gut antibiotic resistome"

**Supplemental tables**

Supplemental Table 1. Categories of foods consumed by subjects for dietary intervention.

|  | Food intervention (number of servings) | | | | | | | |
| --- | --- | --- | --- | --- | --- | --- | --- | --- |
| Sub. ID | Cottage-Cheese | Kefir | Kombucha | Other-drinks | (Probiotic) Shots | Fermented Vegetables | Yoghurt | Total |
| 8004 |  | 19.94 | 11.32 | 18.27 |  | 30.35 | 16.33 | 96.21 |
| 8008 | 1.85 | 1.08 | 2.67 | 8.89 | 6 | 10.66 | 4.45 | 35.6 |
| 8010 |  |  | 22.27 |  |  | 24.81 | 3.14 | 50.22 |
| 8011 |  | 2.66 | 6.65 | 6.53 | 7.62 | 21.92 | 8.42 | 53.8 |
| 8014 |  | 2.66 | 40.7 |  |  |  | 19.05 | 62.41 |
| 8016 | 1.23 | 3.17 | 17.35 | 15.21 |  | 48.58 | 20.74 | 106.28 |
| 8020 |  |  | 58.28 |  |  |  | 23.32 | 81.6 |
| 8021 |  |  | 16.02 | 9.34 | 0.58 | 53.44 | 17.5 | 96.88 |
| 8024 |  | 12 | 31.32 |  |  | 15.83 | 11.99 | 71.14 |
| 8025 |  | 6.73 | 12.91 | 5.16 |  | 15.49 | 12.43 | 52.72 |
| 8026 |  | 11.77 | 19.19 |  |  | 21.84 | 5.59 | 58.39 |
| 8027 |  |  | 41.9 | 5.06 |  | 15.78 | 13.33 | 76.07 |
| 8028 |  | 2.66 | 38.99 |  | 1.35 | 24.23 | 6.95 | 74.18 |
| 8030 |  | 26.64 | 22.43 |  | 24 | 84.71 | 21.93 | 179.71 |
| 8031 | 3 | 3.99 | 2.66 |  | 70 | 23.5 | 21.92 | 125.07 |
| 8032 |  | 11.97 | 22.24 | 6.33 | 33.36 | 1.52 | 5 | 80.42 |
| 8033 |  |  | 9.32 |  |  | 16.02 | 14.84 | 40.18 |
| 8034 | 3.39 | 15.31 | 17.35 | 16.4 |  | 7.6 | 11.47 | 71.52 |
| Total | 9.47 | 120.58 | 393.57 | 91.19 | 142.91 | 416.28 | 238.4 | 1412.4 |

Supplemental STable 2. Fermented food samples assessed in this study.

| Sample # | Description | Year of Purchase | Plate recovered microbiota extraction shotgun | Direct sample extraction shotgun | AR isolate screening | AR isolate illustrated |
| --- | --- | --- | --- | --- | --- | --- |
| Kimchi 1 (K1) | From Korean market A | 2021 | ✔  (mixed) |  |  |  |
| Kimchi 2 (K2) | From Korean restaurant 1 | 2021 | ✔  (mixed) |  |  |  |
| Kimchi 3 (K3) | From Japanese restaurant 2 | 2021 | ✔  (mixed) |  |  |  |
| Kimchi 4 (K4) | From Korean restaurant 3 | 2021 | ✔  (mixed) |  |  |  |
| Kimchi 5 (K5) | From Korean restaurant 3 | 2022 |  |  | ✔ | ✔ |
| Kimchi 6 (K6) | Bottled brand i, from national chain retail store B | 2022 |  |  | ✔ |  |
| Kimchi 7 (K7) | From chain Korean restaurant 4 | 2022 | ✔ | ✔ | ✔ | ✔ |
| Kimchi 8 (K8) | Bottled brand ii, from Asian market C | 2022 |  | ✔ | ✔ | ✔ |
| Kimchi 9 (K9) | Bottled brand iii, from Asian market C | 2022 |  | ✔ | ✔ | ✔ |
| Kimchi 10 (K10) | Bagged brand iv, from national chain retail store D | 2022 |  | ✔ | ✔ | ✔ |
| Kimchi 11 (K11) | From Japanese market E | 2022 |  | ✔ | ✔ | ✔ |
| Kimchi 12 (K12) | Bottled brand v, from Korean Market F | 2022 |  |  | ✔ | ✔ |
| Kimchi 13 (K13) | From Korean Market F | 2022 |  |  | ✔ | ✔ |
| Kimchi 14 (K14) | Bottled brand vi, from national chain retail store G | 2023 |  |  | ✔ | ✔ |
| Cheese 1 (C1) | Blue cheese, pasteurized milk (Germany), from national chain retail store G | 2021 | ✔ (mixed) |  |  |  |
| Cheese 2 (C2) | Blue cheese, raw cow’s milk (Aged > 60 days) (France), from national chain retail store G | 2021 | ✔ (mixed) |  |  |  |
| Cheese 3 (C3) | Blue cheese, pasteurized cow and sheep milk (France), from national chain retail store G | 2021 | ✔ (mixed) |  |  |  |
| Cheese 4 (C4) | Blue cheese, organic pasteurized milk (Unknown), from national chain retail store G | 2021 | ✔ (mixed) |  |  |  |
| Cheese 5 (C5) | Water buffalo milk cheese, pasteurized buffalo milk (Italy), from national chain retail store G | 2022 | ✔ |  | ✔ | ✔ |
| Cheese 6 (C6) | Alpine-style cheese, raw cow’s milk (Aged > 60 days) (United States), from national chain retail store G | 2022 | ✔ |  | ✔ | ✔ |
| Cheese 7 (C7) | Farmhouse table cheese, raw cow’s milk (Aged > 60 days) (United States), from national chain retail store G | 2022 | ✔ |  | ✔ | ✔ |
| Cheese 8 (C8) | Swiss cheese, pasteurized part-skim cow’s milk (Switzerland), from national chain retail store G | 2022 | ✔ |  | ✔ | ✔ |

Supplemental Table 3. Gut antibiotic resistome of subjects with dietary intervention^1^.

|  | Total ARG copies/16S gene | | | | |
| --- | --- | --- | --- | --- | --- |
| Subject ID | "Week-02" | Week0 | Week08 | Week10 | Intervention by |
| 8004 | 0.272186 | 0.284192 | 0.300493 | 0.239926 | Fermented foods |
| 8008 | 0.479228 | 0.508536 | 0.404436 | 0.527567 | Fermented foods |
| 8010 | 0.438165 | 0.460143 | 0.377093 | 0.408678 | Fermented foods |
| 8011 | 0.283978 | 0.241384 |  | 0.345206 | Fermented foods |
| 8014 |  | 0.367376 | 0.420609 | 0.411247 | Fermented foods |
| 8016 | 0.608693 | 0.498886 | 0.611231 | 0.591077 | Fermented foods |
| 8020 | 0.251731 | 0.287428 |  | 0.311607 | Fermented foods |
| 8021 | 0.357738 | 0.428821 | 3.77E-01 | 0.585925 | Fermented foods |
| 8024 | 0.345131 | 0.372126 | 0.364187 | 0.510634 | Fermented foods |
| 8025 | 0.188706 | 0.322113 | 0.308409 | 0.346233 | Fermented foods |
| 8026 | | 0.33126 | 0.515311 | 0.446267 | Fermented foods |
| 8027 | 0.414694 | 0.332016 | 0.455848 | 0.347911 | Fermented foods |
| 8028 | 0.265774 |  |  | 0.351583 | Fermented foods |
| 8030 | 0.407724 | 0.514995 | 0.598983 | 0.448453 | Fermented foods |
| 8032 | 0.383879 | 0.416004 | 0.468486 | 0.718217 | Fermented foods |
| 8033 | 0.290082 | 0.272963 | 0.236222 | 0.23334 | Fermented foods |
| 8034 | 0.297562 | 0.364338 | 0.363455 | 0.287874 | Fermented foods |
| 8001 |  | 0.376419 | 0.316027 | 0.310167 | Plant fibers |
| 8002 | 0.274209 | 0.289648 | 0.264493 | 0.265522 | Plant fibers |
| 8003 | 0.624062 | 0.481308 | 0.447535 | 0.394988 | Plant fibers |
| 8006 | 0.219233 | 0.219967 | 0.233956 | 0.253023 | Plant fibers |
| 8009 | 0.182759 | 0.449417 | 0.415087 | 0.433871 | Plant fibers |
| 8017 | 0.765387 | 0.246743 |  | 0.342993 | Plant fibers |
| 8018 | 0.259782 |  | 0.241797 | 0.279882 | Plant fibers |
| 8022 | 0.364975 |  | 0.534551 | 0.44805 | Plant fibers |
| 8023 |  | 0.467738 | 0.385584 | 0.507301 | Plant fibers |
| 8029 | 0.397652 | 0.313326 | 0.326775 | 0.281955 | Plant fibers |
| 8036 | 0.381629 | 0.421726 |  | 0.45393 | Plant fibers |
| 8037 | 0.354639 | 0.320164 | 0.283361 | 0.384654 | Plant fibers |
| 8038 | 0.338513 | 0.269464 | 0.420975 | 0.311128 | Plant fibers |
| 8039 | 0.409468 | 0.552285 | 0.551152 | 0.555141 | Plant fibers |
| 8041 | 0.499627 | 0.501698 | 0.510558 | 0.526228 | Plant fibers |

^1^Numerical readings in total ARG copies per 16S gene.

Supplemental Table 4. Top 5 genera classified of microbiota recovered from BHI agar plates of pooled 4 kimchi samples purchased in 2021.

| Sample | Top genera | Percentage |
| --- | --- | --- |
| Mix_Kimchi_BHI | *Bacillus* | 58.29% |
|  | *Serratia* | 13.42% |
|  | *Pseudomonas* | 12.11% |
|  | *Leuconostoc* | 9.92% |
|  | Other | 4.72% |
|  | *Lelliottia* | 1.55% |
| Mix_Kimchi_BHI_amp | *Pseudomonas* | 46.63% |
|  | *Bacillus* | 41.71% |
|  | *Brucella* | 5.92% |
|  | *Enterobacter* | 2.71% |
|  | Other | 1.70% |
|  | *Rahnella* | 1.34% |
| Mix_Kimchi_BHI_tet | *Latilactobacillus* | 34.33% |
|  | *Bacillus* | 33.42% |
|  | *Serratia* | 18.88% |
|  | *Stenotrophomonas* | 5.65% |
|  | *Lactococcus* | 4.14% |
|  | Other | 3.57% |

Supplemental Table 5: Top 5 genera classified of microbiota recovered from BHI agar plates of pooled 4 cheese samples purchased in 2021.

| Sample | Top genera | Percentage |
| --- | --- | --- |
| Mix_Cheese_BHI | *Lactococcus* | 47.32% |
|  | *Lacticaseibacillus* | 25.26% |
|  | *Staphylococcus* | 23.19% |
|  | Other | 2.04% |
|  | *Enterococcus* | 1.09% |
|  | *Streptococcus* | 1.09% |
| Mix_Cheese_BHI_tet | *Staphylococcus* | 52.84% |
|  | *Bacillus* | 31.53% |
|  | *Streptococcus* | 12.24% |
|  | *Microbacterium* | 1.92% |
|  | Other | 1.29% |
|  | *Bacteroides* | 0.17% |

Supplemental Table 6: Top 5 genera classified of BHI-recovered microbiota of individual cheese samples purchased in 2022.

| Sample | Top genera | Percentage |
| --- | --- | --- |
| Cheese5-BHI | *Mammaliicoccus* | 64.41% |
|  | *Staphylococcus* | 18.62% |
|  | *Carnobacterium* | 5.26% |
|  | Other | 4.49% |
|  | *Glutamicibacter* | 3.73% |
|  | *Psychrobacter* | 3.51% |
| Cheese5-BHI-amp | Other | 68.49% |
|  | *Stenotrophomonas* | 9.89% |
|  | *Pseudomonas* | 7.20% |
|  | *Burkholderia* | 5.91% |
|  | *Alcanivorax* | 5.31% |
|  | *Streptococcus* | 3.19% |
| Cheese5-BHI-tet | Other | 64.56% |
|  | *Stenotrophomonas* | 11.03% |
|  | *Pseudomonas* | 7.46% |
|  | *Burkholderia* | 6.17% |
|  | *Alcanivorax* | 5.66% |
|  | *Streptococcus* | 5.14% |
| Cheese6-BHI | *Staphylococcus* | 64.68% |
|  | *Enterococcus* | 22.47% |
|  | *Lacticaseibacillus* | 12.22% |
|  | Other | 0.29% |
|  | *Streptococcus* | 0.28% |
|  | *Leuconostoc* | 0.07% |
| Cheese7-BHI | *Staphylococcus* | 73.00% |
|  | *Lactococcus* | 26.05% |
|  | *Leuconostoc* | 0.40% |
|  | Other | 0.32% |
|  | *Lacticaseibacillus* | 0.13% |
|  | *Streptococcus* | 0.11% |
| Cheese7-BHI-amp | Other | 50.11% |
|  | *Pseudomonas* | 30.32% |
|  | *Burkholderia* | 5.63% |
|  | *Staphylococcus* | 5.18% |
|  | *Streptococcus* | 5.16% |
|  | *Leuconostoc* | 3.60% |
| Cheese7-BHI-tet | *Staphylococcus* | 99.71% |
|  | Other | 0.10% |
|  | *Enterococcus* | 0.09% |
|  | *Leuconostoc* | 0.05% |
|  | *Mammaliicoccus* | 0.03% |
|  | *Lactococcus* | 0.02% |
| Cheese8-BHI | *Enterococcus* | 44.89% |
|  | *Tetragenococcus* | 16.18% |
|  | *Staphylococcus* | 13.08% |
|  | *Lactococcus* | 12.68% |
|  | Other | 11.26% |
|  | *Carnobacterium* | 1.91% |
