## Supplemental table 7 and 8 for "High prevalence of antibiotic resistance in traditionally fermented foods as a critical risk factor for host gut antibiotic resistome"

Supplementary Table 7. Sensititre MIC for representative Gram-negative isolates from kimchi*.

| Drug (Test Range, µg/mL) | *Kleb pneumoniae*  K7-6 | *Rahnella aquatilis*  K11-3 | *Serratia marcescens*  K13-1 |
| --- | --- | --- | --- |
| Amikacin (8-64) | **16** | <8 | 8 |
| *Ampicillin* (4-32) | **>32** | **>32** | **>32** |
| Ampicillin/sulbactam  2:1 ratio (4/2-32/16) | 8/4 | <4/2 | **>32/16** |
| Aztreonam (4-32) | <4 | <4 | **16** |
| *Cefazolin* (4-32) | 4 | **>32** | **>32** |
| *Cefepime* (4-32) | <4 | <4 | 4 |
| *Cephalothin* (2-16) | 4 | **>16** | **>16** |
| *Meropenem* (1-8) | <1 | <1 | **4** |
| *Ertapenem* (2-16) | <2 | <2 | **4** |
| *Cefuroxime* (4-32) | <4 | **>32** | **>32** |
| Gentamicin (2-16) | 4 | <2 | **>16** |
| Ciprofloxacin (0.5-4) | <0.5 | <0.5 | <0.5 |
| *Piperacillin*/tazobactam constant 4 (16/4-128/4) | <16/4 | <16/4 | **>128/4** |
| *Cefoxitin* (4-32) | 8 | <4 | **>32** |
| Trimethoprim/sulfamethoxazole (0.5/9.5-4/76) | <0.5/9.5 | <0.5/9.5 | **>4/76** |
| *Cefpodoxime* (2-16) | <2 | **16** | 8 |
| *Ceftazidime* (1-32) | <1 | <1 | **>32** |
| Tobramycin (4-8) | <4 | <4 | >8 |
| Tigecycline (1-8) | <1 | <1 | 1 |
| Ticarcillin/clavulanic constant 2 (16/2-64/2) | <16/2 | <16/2 | 16/2 |
| *Ceftriaxone* (1-64) | <1 | **8** | **>64** |
| Tetracycline (0.5-16) | 2 | 4 | **>16** |

*Bold MIC numbers: strongly indicative of the isolates being resistant to the corresponding antibiotics based on references (CLSI, 2020; Marchaim, 2014; Papich & Lindeman, 2018). *Italicized antibiotics:* penicillin and derivatives. *Italicized and highlighted antibiotics:* 4^th^ generation of **cephalosporin antibiotic (***Cefepime)* **or** Carbapenem antibiotics (*Meropenem, Ertapenem).*

Supplementary Table 8. Sensititre MIC for representative Gram-positive isolates from kimchi and artisan cheeses*.

| Drug (Test Range, ug/mL) | *Weissella* spp.  K5-3 | *Lb. brevis* K10-1 | *Leuc. Citreum*  K12-1 | *Lb. brevis*  K14-2 | *Staphy xylosus*  C6-2 | *E. faecalis*  C6-11 |
| --- | --- | --- | --- | --- | --- | --- |
| Erythromycin (0.25-4) | **>4** | <0.25 | <0.25 | <0.25 | 0.5 | **>4** |
| Clindamycin (0.12-2) | >2 | 2 | <0.12 | <0.12 | 0.25 | >2 |
| Quinupristin/ dalfopristin (0.12-4) | >4 | 1 | >0.5 | 0.5 | 0.5 | >4 |
| Daptomycin (0.25-8) | **>8** | **>8** | >0.25 | **>8** | 2 | **>8** |
| Vancomycin (1-128) | **>128** | **>128** | **>128** | **>128** | 1 | **>128** |
| Tetracycline (2-16) | **>16** | **16** | **>16** | **16** | **>16** | <2 |
| *Ampicillin* (0.12-16) | **>16** | 2 | >0.5 | 2 | 0.25 | 1 |
| Gentamicin (2-16/500) | **>500** | <2 | >8 | 2 | <2 | **>16** |
| Levofloxacin (0.25-8) | **>8** | >2 | >1 | 4 | 0.5 | 4 |
| Linezolid (0.5-8) | **>8** | 4 | >2 | 4 | 2 | 4 |
| *Ceftriaxone* (3^rd^) (8-64) | **>64** | **>64** | <8 | **>64** | 8 | **>64** |
| Streptomycin (1000) | **>1000** | <1000 | <1000 | <1000 | <1000 | <1000 |
| *Penicillin* (0.06-8) | **>8** | **8** | 0.25 | 4 | 0.25 | 1 |
| Rifampin (0.5-4) | **>4** | 0.5 | >1 | 0.5 | <0.5 | 4 |
| Gatifloxacin (1-8) | **>8** | 1 | <1 | 1 | <1 | 1 |
| Ciprofloxacin (0.5-2) | **>2** | **>2** | **>2** | **>2** | 0.5 | **>2** |
| Trimethoprim/ sulfamethoxazole (0.5/9.5-4/76) | **>4/76** | **>4/76** | **>4/76** | **>4/76** | <0.5/9.5 | **>4/76** |
| *Oxacillin* + 2%NaCl (0.25-8) | **>8** | **>8** | **>1** | **>8** | **1** | **>8** |

*Bold MIC numbers: strongly indicative of the isolates being resistant to the corresponding antibiotics based on references (CLSI, 2020). *Italicized antibiotics:* penicillin and derivatives.
