## Supplemental Table 9 resistome for "High prevalence of antibiotic resistance in traditionally fermented foods as a critical risk factor for host gut antibiotic resistome"

Supplemental Table 9. Fecal antibiotic resistome of subjects by fermented foods intervention.

| ARG_types | 8004_B2W | 8004_0W | 8004_8W | 8004_10W | 8008_B2W | 8008_0W |
|---|---|---|---|---|---|---|
| aminoglycoside | 0.049838 | 0.017378 | 0.086563 | 0.020008 | 0.038533 | 0.060416 |
| bacitracin | 0.012912 | 0.013694 | 0.015595 | 0.027826 | 0.025612 | 0.030658 |
| beta-lactam | 0.036763 | 0.016648 | 0.027417 | 0.016897 | 0.064398 | 0.040065 |
| bleomycin | 0 | 0 | 0 | 0 | 0 | 0 |
| carbomycin | 0 | 0 | 0 | 0 | 0 | 0 |
| chloramphenicol | 0.000317 | 0.003902 | 0.002014 | 0.001034 | 0.001264 | 0.001562 |
| fosfomycin | 9.92E-05 | 0 | 0 | 0 | 2.91E-05 | 0 |
| fosmidomycin | 0.002273 | 0.003419 | 0.000821 | 0.000453 | 0.000995 | 0.000402 |
| fusaric-acid | 0 | 0 | 0 | 0 | 0 | 0 |
| fusidic-acid | 0 | 0 | 0 | 0 | 0 | 0 |
| kasugamycin | 0 | 0 | 0 | 0 | 5.24E-05 | 0 |
| macrolide-lincosamide-streptogramin | 0.01064 | 0.027375 | 0.012997 | 0.008727 | 0.109551 | 0.149931 |
| multidrug | 0.005429 | 0.004625 | 0.006106 | 0.00522 | 0.005858 | 0.004065 |
| polymyxin | 0.000472 | 0 | 0.000152 | 2.83E-05 | 7.00E-05 | 0.000164 |
| puromycin | 0 | 0 | 0 | 0 | 0 | 0 |
| quinolone | 0.000331 | 0.001356 | 0.000486 | 0.000254 | 0 | 0 |
| rifamycin | 0 | 8.30E-05 | 0 | 0 | 3.29E-05 | 0 |
| spectinomycin | 0 | 0 | 0 | 0 | 0 | 0 |
| sulfonamide | 0.000109 | 0 | 0 | 1.23E-05 | 0.000346 | 0 |
| tetracenomycin_C | 0 | 0 | 0 | 0 | 0 | 0 |
| tetracycline | 0.122468 | 0.153039 | 0.099887 | 0.089917 | 0.209427 | 0.192851 |
| trimethoprim | 0 | 0 | 0 | 0 | 0 | 0 |
| unclassified | 0.000373 | 0.000402 | 0.000366 | 0.000373 | 0.002036 | 0.000817 |
| vancomycin | 0.030161 | 0.042271 | 0.048089 | 0.069177 | 0.021023 | 0.027604 |
| SUM | 0.272186 | 0.284192 | 0.300493 | 0.239926 | 0.479228 | 0.508536 |

| ARG_types | 8008_8W | 8008_10W | 8010_B2W | 8010_0W | 8010_8W | 8010_10W |
|---|---|---|---|---|---|---|
| aminoglycoside | 0.040362 | 0.042148 | 0.006121 | 0.004569 | 0.007132 | 0.006964 |
| bacitracin | 0.013811 | 0.01755 | 0.036304 | 0.040535 | 0.017888 | 0.02588 |
| beta-lactam | 0.068901 | 0.064998 | 0.087796 | 0.109024 | 0.087224 | 0.089338 |
| bleomycin | 0 | 0 | 0 | 0 | 0 | 0 |
| carbomycin | 0 | 0 | 0 | 0 | 0 | 0 |
| chloramphenicol | 0.000253 | 0.001852 | 0.009375 | 0.008305 | 0.007467 | 0.008798 |
| fosfomycin | 0 | 0 | 0 | 0.000615 | 0 | 0 |
| fosmidomycin | 0.000413 | 0.000916 | 0.000686 | 0.000595 | 0.000538 | 0.000627 |
| fusaric-acid | 0 | 0 | 0 | 0 | 0 | 0 |
| fusidic-acid | 0 | 0 | 0 | 0 | 0 | 0 |
| kasugamycin | 4.88E-05 | 0.000485 | 0.000396 | 0.00079 | 0.000103 | 0.000237 |
| macrolide-lincosamide-streptogramin | 0.08552 | 0.119767 | 0.062919 | 0.068587 | 0.06878 | 0.056612 |
| multidrug | 0.003398 | 0.010453 | 0.006997 | 0.008552 | 0.00425 | 0.00413 |
| polymyxin | 0 | 0.000236 | 0.000222 | 0.000116 | 0 | 0 |
| puromycin | 0 | 0 | 0 | 0 | 0 | 0 |
| quinolone | 0 | 0 | 0.000199 | 0.000105 | 0.000236 | 0.000111 |
| rifamycin | 0 | 2.70E-05 | 0 | 0 | 0 | 4.66E-05 |
| spectinomycin | 0 | 0 | 0 | 0 | 0 | 0 |
| sulfonamide | 0 | 0.00064 | 0 | 2.85E-05 | 0 | 0 |
| tetracenomycin_C | 0 | 0 | 0 | 0 | 0 | 0 |
| tetracycline | 0.179194 | 0.238526 | 0.214198 | 0.206891 | 0.175714 | 0.205469 |
| trimethoprim | 0 | 0 | 0 | 0 | 0 | 0 |
| unclassified | 0.000801 | 0.002606 | 0.002091 | 0.002568 | 0.000338 | 0.001194 |
| vancomycin | 0.011734 | 0.027363 | 0.010859 | 0.008862 | 0.007424 | 0.009274 |
| SUM | 0.404436 | 0.527567 | 0.438165 | 0.460143 | 0.377093 | 0.408678 |

| ARG_types | 8011_B2W | 8011_0W | 8011_10W | 8016_B2W | 8016_0W | 8016_8W |
|---|---|---|---|---|---|---|
| aminoglycoside | 0.008647 | 0.006303 | 0.010437 | 0.022229 | 0.020013 | 0.034928 |
| bacitracin | 0.031929 | 0.034485 | 0.020079 | 0.0215 | 0.024947 | 0.011508 |
| beta-lactam | 0.066213 | 0.044235 | 0.088566 | 0.005315 | 0.00599 | 0.030814 |
| bleomycin | 0 | 0 | 0 | 0 | 0 | 0 |
| carbomycin | 0 | 0 | 0 | 0 | 0 | 0 |
| chloramphenicol | 0.000462 | 0.003894 | 0.000153 | 0.002805 | 0.004325 | 0.007127 |
| fosfomycin | 0 | 9.94E-05 | 0 | 0 | 0 | 0 |
| fosmidomycin | 0.002102 | 0.003026 | 0.002005 | 0.002285 | 0.001261 | 0.002698 |
| fusaric-acid | 0 | 0 | 0 | 0 | 0 | 0 |
| fusidic-acid | 0 | 0 | 0 | 0 | 0 | 0 |
| kasugamycin | 8.76E-05 | 0.000729 | 0.000103 | 0 | 0 | 0 |
| macrolide-lincosamide-streptogramin | 0.017156 | 0.013175 | 0.022469 | 0.062833 | 0.042229 | 0.073272 |
| multidrug | 0.005641 | 0.01655 | 0.005736 | 0.006647 | 0.005067 | 0.010103 |
| polymyxin | 0.000134 | 0.000594 | 2.26E-05 | 2.25E-05 | 0.0001 | 1.76E-05 |
| puromycin | 0 | 0 | 0 | 0 | 0 | 0 |
| quinolone | 0 | 7.01E-05 | 0 | 0 | 0 | 0 |
| rifamycin | 0 | 2.38E-05 | 0 | 0 | 0.000131 | 0 |
| spectinomycin | 0 | 0 | 0 | 0 | 0 | 0 |
| sulfonamide | 0 | 0 | 0 | 0 | 0.000111 | 0.000128 |
| tetracenomycin_C | 0 | 0 | 0 | 0 | 0 | 0 |
| tetracycline | 0.140482 | 0.103002 | 0.178694 | 0.45042 | 0.366827 | 0.404997 |
| trimethoprim | 0 | 0 | 4.90E-05 | 0 | 0 | 7.56E-05 |
| unclassified | 0.000932 | 0.002969 | 0.00076 | 0.000448 | 0.000282 | 0.000979 |
| vancomycin | 0.010193 | 0.01223 | 0.016133 | 0.034187 | 0.027604 | 0.034582 |
| SUM | 0.283978 | 0.241384 | 0.345206 | 0.608693 | 0.498886 | 0.611231 |

| ARG_types | 8016_10W | 8020_B2W | 8020_0W | 8020_10W | 8021_B2W | 8021_0W |
|---|---|---|---|---|---|---|
| aminoglycoside | 0.053946 | 0.0088 | 0.008774 | 0.010426 | 0.025025 | 0.02431 |
| bacitracin | 0.018131 | 0.026811 | 0.030107 | 0.021996 | 0.018753 | 0.015698 |
| beta-lactam | 0.008353 | 0.039809 | 0.035977 | 0.042815 | 0.061037 | 0.083868 |
| bleomycin | 0 | 0 | 0 | 0 | 0 | 0 |
| carbomycin | 0 | 0 | 0 | 0 | 0 | 0 |
| chloramphenicol | 0.004622 | 0.00034 | 0.001459 | 0.000822 | 7.16E-05 | 0.000216 |
| fosfomycin | 0 | 0 | 0 | 0 | 0 | 5.73E-05 |
| fosmidomycin | 0.001358 | 0.001116 | 0.003022 | 0.000354 | 0.001615 | 0.001389 |
| fusaric-acid | 0 | 0 | 0 | 0 | 0 | 0 |
| fusidic-acid | 0 | 0 | 0 | 0 | 0 | 0 |
| kasugamycin | 0 | 0 | 0.001047 | 0.00013 | 7.75E-05 | 0.00011 |
| macrolide-lincosamide-streptogramin | 0.049231 | 0.007267 | 0.011724 | 0.008227 | 0.039288 | 0.054158 |
| multidrug | 0.00893 | 0.005118 | 0.027287 | 0.006476 | 0.005734 | 0.006474 |
| polymyxin | 0.000102 | 0 | 0.000762 | 4.88E-05 | 5.34E-05 | 0.000201 |
| puromycin | 0 | 0 | 0 | 0 | 0 | 0 |
| quinolone | 0 | 0 | 0 | 0 | 0.000114 | 0.000112 |
| rifamycin | 5.73E-05 | 0 | 0 | 0 | 0 | 0 |
| spectinomycin | 0 | 0 | 0 | 0 | 0 | 0 |
| sulfonamide | 0.000186 | 0 | 0 | 0 | 0.048849 | 0.045477 |
| tetracenomycin_C | 0 | 0 | 0 | 0 | 0 | 0 |
| tetracycline | 0.408171 | 0.156819 | 0.154451 | 0.213953 | 0.141041 | 0.17557 |
| trimethoprim | 0 | 0 | 0 | 0 | 0 | 0 |
| unclassified | 0.001345 | 0.001396 | 0.007383 | 0.001435 | 0.000741 | 0.00117 |
| vancomycin | 0.036644 | 0.004258 | 0.005436 | 0.004924 | 0.015338 | 0.02001 |
| SUM | 0.591077 | 0.251731 | 0.287428 | 0.311607 | 0.357738 | 0.428821 |

| ARG_types | 8021_8W | 8021_10W | 8024_B2W | 8024_0W | 8024_8W | 8024_10W |
|---|---|---|---|---|---|---|
| aminoglycoside | 0.021485 | 0.119437 | 0.030749 | 0.009134 | 0.017049 | 0.029256 |
| bacitracin | 0.021255 | 0.018636 | 0.011112 | 0.0205 | 0.017149 | 0.014313 |
| beta-lactam | 0.036809 | 0.088092 | 0.051006 | 0.073257 | 0.066259 | 0.039388 |
| bleomycin | 0 | 0 | 0 | 0 | 0 | 0 |
| carbomycin | 0 | 0 | 0 | 0 | 0 | 0 |
| chloramphenicol | 0 | 0.000184 | 0.000169 | 0.000108 | 0.008072 | 0.014404 |
| fosfomycin | 0 | 0 | 0.000152 | 1.85E-05 | 0 | 0 |
| fosmidomycin | 0.00162 | 0.000883 | 0.001456 | 0.001215 | 0.001681 | 0.004283 |
| fusaric-acid | 0 | 0 | 0 | 0 | 0 | 0 |
| fusidic-acid | 0 | 0 | 0 | 0 | 0 | 0 |
| kasugamycin | 8.50E-05 | 0.000141 | 0.000389 | 0.000602 | 0 | 0 |
| macrolide-lincosamide-streptogramin | 0.058782 | 0.050691 | 0.046875 | 0.050701 | 0.06759 | 0.089968 |
| multidrug | 0.005931 | 0.005505 | 0.02187 | 0.016356 | 0.008626 | 0.006363 |
| polymyxin | 0.000141 | 3.08E-05 | 0.000474 | 0.000548 | 0.000274 | 0.00055 |
| puromycin | 0 | 0 | 0 | 0 | 0 | 0 |
| quinolone | 0.000122 | 2.16E-05 | 0.000201 | 0.000194 | 0 | 0 |
| rifamycin | 0 | 0 | 0 | 0 | 0 | 0 |
| spectinomycin | 0 | 0 | 0 | 0 | 0 | 0 |
| sulfonamide | 0.011035 | 0.104362 | 0.000192 | 0.000509 | 0 | 0 |
| tetracenomycin_C | 0 | 0 | 0 | 0 | 0 | 0 |
| tetracycline | 0.186868 | 0.181596 | 0.16549 | 0.179228 | 0.16759 | 0.295996 |
| trimethoprim | 0 | 0 | 0.000149 | 0.000376 | 0 | 0.000163 |
| unclassified | 0.001159 | 0.001085 | 0.002372 | 0.003462 | 0.000585 | 0.000289 |
| vancomycin | 0.031303 | 0.015261 | 0.012475 | 0.015917 | 0.009311 | 0.015662 |
| SUM | 0.376594 | 0.585925 | 0.345131 | 0.372126 | 0.364187 | 0.510634 |

| ARG_types | 8025_B2W | 8025_0W | 8025_8W | 8025_10W | 8027_B2W | 8027_0W |
|---|---|---|---|---|---|---|
| aminoglycoside | 0.013618 | 0.019032 | 0.017864 | 0.018384 | 0.011315 | 0.011032 |
| bacitracin | 0.020335 | 0.013422 | 0.009863 | 0.011148 | 0.023201 | 0.029217 |
| beta-lactam | 0.00459 | 0.026535 | 0.02314 | 0.030871 | 0.082703 | 0.050974 |
| bleomycin | 0 | 0 | 0 | 0 | 0 | 0 |
| carbomycin | 0 | 0 | 1.11E-05 | 0 | 0 | 0 |
| chloramphenicol | 0.000262 | 0.000222 | 0 | 0.000367 | 0.014278 | 0.011503 |
| fosfomycin | 0 | 0 | 0 | 0 | 0 | 0.000112 |
| fosmidomycin | 0.000435 | 0.001145 | 0.001618 | 0.001573 | 0.00287 | 0.000776 |
| fusaric-acid | 0 | 0 | 0 | 0 | 0 | 0 |
| fusidic-acid | 0 | 0 | 0 | 0 | 0 | 0 |
| kasugamycin | 0 | 0.000169 | 3.59E-05 | 0.000159 | 0.000952 | 0.000564 |
| macrolide-lincosamide-streptogramin | 0.005457 | 0.006331 | 0.007463 | 0.009449 | 0.043744 | 0.029784 |
| multidrug | 0.004824 | 0.010283 | 0.008341 | 0.015062 | 0.031113 | 0.020926 |
| polymyxin | 4.19E-05 | 0.000617 | 0.000483 | 0.000617 | 0.00059 | 0.000601 |
| puromycin | 0 | 0 | 0 | 0 | 0 | 0 |
| quinolone | 0 | 0 | 0 | 0 | 0 | 0 |
| rifamycin | 2.59E-05 | 0 | 0 | 0 | 0 | 0 |
| spectinomycin | 0 | 0 | 0 | 0 | 0 | 0 |
| sulfonamide | 0 | 0 | 4.56E-05 | 2.85E-05 | 0 | 0 |
| tetracenomycin_C | 0 | 0 | 0 | 0 | 0 | 0 |
| tetracycline | 0.120352 | 0.226278 | 0.223252 | 0.240404 | 0.186418 | 0.152915 |
| trimethoprim | 0 | 0 | 0 | 0 | 0 | 0 |
| unclassified | 0.000249 | 0.000602 | 0.000716 | 0.001586 | 0.004621 | 0.004196 |
| vancomycin | 0.018515 | 0.017478 | 0.015577 | 0.016587 | 0.012887 | 0.019417 |
| SUM | 0.188706 | 0.322113 | 0.308409 | 0.346233 | 0.414694 | 0.332016 |

| ARG_types | 8027_8W | 8027_10W | 8028_B2W | 8028_10W | 8030_B2W | 8030_0W |
|---|---|---|---|---|---|---|
| aminoglycoside | 0.007004 | 0.001977 | 0.006825 | 0.003313 | 0.003957 | 0.003492 |
| bacitracin | 0.025933 | 0.039349 | 0.028681 | 0.019041 | 0.013653 | 0.018758 |
| beta-lactam | 0.082694 | 0.09616 | 0.022662 | 0.044527 | 0.041795 | 0.059637 |
| bleomycin | 0 | 0 | 0 | 0 | 0 | 0 |
| carbomycin | 0 | 0 | 0 | 0 | 0 | 0 |
| chloramphenicol | 0.032713 | 0.007541 | 0.001417 | 0.001121 | 0.002984 | 0.002794 |
| fosfomycin | 7.68E-05 | 2.62E-05 | 0 | 0 | 0 | 0 |
| fosmidomycin | 0.001422 | 0.000762 | 0.0012 | 0.001237 | 0.003467 | 0.00587 |
| fusaric-acid | 0 | 0 | 0 | 0 | 0 | 0 |
| fusidic-acid | 0 | 0 | 0 | 0 | 0 | 0 |
| kasugamycin | 0 | 0.000267 | 0.000297 | 0.00027 | 0.001849 | 0.003571 |
| macrolide-lincosamide-streptogramin | 0.062988 | 0.033243 | 0.023572 | 0.043125 | 0.007301 | 0.014773 |
| multidrug | 0.007413 | 0.013701 | 0.008387 | 0.003928 | 0.046509 | 0.099948 |
| polymyxin | 0.000111 | 0.000225 | 9.65E-05 | 2.44E-05 | 0.00164 | 0.003446 |
| puromycin | 0 | 0 | 0 | 0 | 0 | 0 |
| quinolone | 0 | 0 | 0 | 0 | 0 | 0 |
| rifamycin | 0.000225 | 0 | 3.71E-05 | 2.29E-05 | 0 | 0 |
| spectinomycin | 0 | 0 | 0 | 0 | 0 | 0 |
| sulfonamide | 0 | 0 | 0.000362 | 5.69E-05 | 0 | 0 |
| tetracenomycin_C | 0 | 0 | 0 | 0 | 0 | 0 |
| tetracycline | 0.219362 | 0.142647 | 0.139937 | 0.218284 | 0.258655 | 0.269378 |
| trimethoprim | 0 | 0 | 0.000178 | 2.11E-05 | 0 | 0 |
| unclassified | 0.00019 | 0.001852 | 0.001706 | 0.000833 | 0.011789 | 0.023284 |
| vancomycin | 0.015714 | 0.01016 | 0.030414 | 0.015781 | 0.014125 | 0.010043 |
| SUM | 0.455848 | 0.347911 | 0.265774 | 0.351583 | 0.407724 | 0.514995 |

| ARG_types | 8030_8W | 8030_10W | 8032_B2W | 8032_0W | 80326 | 8032_10W |
|---|---|---|---|---|---|---|
| aminoglycoside | 0.004206 | 0.005336 | 0.010154 | 0.012352 | 0.01859 | 0.020669 |
| bacitracin | 0.021754 | 0.018914 | 0.030424 | 0.02973 | 0.020186 | 0.019895 |
| beta-lactam | 0.064547 | 0.04331 | 0.108333 | 0.104187 | 0.112423 | 0.194018 |
| bleomycin | 0 | 0 | 0 | 0 | 0 | 0 |
| carbomycin | 0 | 0 | 0 | 0 | 0 | 0 |
| chloramphenicol | 0.004356 | 0.00638 | 0.000485 | 0.000443 | 0.0005 | 0.001759 |
| fosfomycin | 0 | 0 | 0 | 0 | 0 | 0 |
| fosmidomycin | 0.006871 | 0.003084 | 0.00255 | 0.003267 | 0.00387 | 0.001717 |
| fusaric-acid | 0 | 0 | 0 | 0 | 0 | 0 |
| fusidic-acid | 0 | 0 | 0 | 0 | 0 | 0 |
| kasugamycin | 0.005312 | 0.001134 | 1.43E-05 | 0.000317 | 0.000174 | 0.000237 |
| macrolide-lincosamide-streptogramin | 0.011741 | 0.006792 | 0.037566 | 0.045211 | 0.080362 | 0.140909 |
| multidrug | 0.128124 | 0.030072 | 0.008182 | 0.0122 | 0.01051 | 0.007857 |
| polymyxin | 0.004743 | 0.001156 | 0.000191 | 0.000384 | 0.000131 | 0.000292 |
| puromycin | 0 | 0 | 0 | 0 | 0 | 0 |
| quinolone | 0 | 0 | 0 | 0 | 0 | 0 |
| rifamycin | 0 | 0 | 0 | 0 | 0 | 0 |
| spectinomycin | 0 | 0 | 0 | 0 | 0 | 0 |
| sulfonamide | 0 | 3.91E-05 | 0.000196 | 0.000548 | 0.000166 | 0.000414 |
| tetracenomycin_C | 0 | 0 | 0 | 0 | 0 | 0 |
| tetracycline | 0.304266 | 0.307309 | 0.15508 | 0.183219 | 0.206756 | 0.312681 |
| trimethoprim | 0 | 0 | 0.00037 | 0.000435 | 0 | 0 |
| unclassified | 0.031021 | 0.006699 | 0.001186 | 0.001832 | 0.001535 | 0.001196 |
| vancomycin | 0.012044 | 0.018228 | 0.029148 | 0.021879 | 0.013282 | 0.016573 |
| SUM | 0.598983 | 0.448453 | 0.383879 | 0.416004 | 0.468486 | 0.718217 |

| ARG_types | 8033_B2W | 8033_0W | 8033_8W | 8033_10W | 8034_B2W | 8034_0W |
|---|---|---|---|---|---|---|
| aminoglycoside | 0.022823 | 0.038818 | 0.023776 | 0.032099 | 0.086401 | 0.114621 |
| bacitracin | 0.021668 | 0.026406 | 0.019924 | 0.034706 | 0.018141 | 0.012035 |
| beta-lactam | 0.014785 | 0.007618 | 0.009444 | 0.006796 | 0.00778 | 0.028169 |
| bleomycin | 0 | 0 | 0 | 0 | 0 | 0 |
| carbomycin | 0 | 0 | 0 | 0 | 0 | 0 |
| chloramphenicol | 0.001603 | 0.002098 | 0.000584 | 0.000699 | 0.006226 | 0.003318 |
| fosfomycin | 0 | 3.39E-05 | 7.96E-05 | 9.60E-05 | 0 | 0 |
| fosmidomycin | 0.001637 | 0.000402 | 0.000615 | 0.000424 | 0.000335 | 0.000723 |
| fusaric-acid | 0 | 0 | 0 | 0 | 0 | 0 |
| fusidic-acid | 0 | 0 | 0 | 0 | 0 | 0 |
| kasugamycin | 0.001308 | 0.000203 | 0 | 2.93E-05 | 0 | 0.000108 |
| macrolide-lincosamide-streptogramin | 0.019493 | 0.010098 | 0.00802 | 0.006336 | 0.04634 | 0.067716 |
| multidrug | 0.025762 | 0.008855 | 0.007294 | 0.00531 | 0.011028 | 0.007717 |
| polymyxin | 0.000811 | 0.00036 | 0.000365 | 3.38E-05 | 0 | 8.97E-05 |
| puromycin | 0 | 0 | 0 | 0 | 0 | 0 |
| quinolone | 0 | 0 | 0 | 0 | 0 | 0 |
| rifamycin | 0 | 0 | 0 | 0 | 0 | 0 |
| spectinomycin | 0 | 0 | 0 | 0 | 0 | 0 |
| sulfonamide | 0.001625 | 0.001696 | 0.001875 | 0.001199 | 0 | 0 |
| tetracenomycin_C | 0 | 0 | 0 | 0 | 0 | 0 |
| tetracycline | 0.163946 | 0.150063 | 0.131815 | 0.118135 | 0.103561 | 0.114939 |
| trimethoprim | 0 | 0 | 0 | 0 | 0 | 0 |
| unclassified | 0.00567 | 0.001559 | 0.001332 | 0.000296 | 0.000425 | 0.000598 |
| vancomycin | 0.008951 | 0.024752 | 0.031098 | 0.027179 | 0.017324 | 0.014304 |
| SUM | 0.290082 | 0.272963 | 0.236222 | 0.23334 | 0.297562 | 0.364338 |

| ARG_types | 8034_8W | 8034_10W | 8026_0W | 8026_8W | 8026_10W | 8014_0W |
|---|---|---|---|---|---|---|
| aminoglycoside | 0.114143 | 0.074065 | 0.003999 | 0.00514 | 0.009192 | 0.013831 |
| bacitracin | 0.01322 | 0.011101 | 0.023598 | 0.021598 | 0.016883 | 0.02 |
| beta-lactam | 0.017947 | 0.010226 | 0.075633 | 0.252866 | 0.183279 | 0.029829 |
| bleomycin | 0 | 0 | 0 | 0 | 0 | 0 |
| carbomycin | 0 | 0 | 0 | 0 | 0 | 0 |
| chloramphenicol | 0.006683 | 0.006137 | 0.004355 | 0.005086 | 0.003395 | 0.001109 |
| fosfomycin | 0 | 0 | 0 | 0 | 0 | 0 |
| fosmidomycin | 0.000358 | 0.000399 | 1.67E-05 | 5.25E-05 | 1.92E-05 | 0.001173 |
| fusaric-acid | 0 | 0 | 0 | 0 | 0 | 0 |
| fusidic-acid | 0 | 0 | 0 | 0 | 0 | 0 |
| kasugamycin | 1.42E-05 | 0 | 0 | 0 | 1.71E-05 | 0 |
| macrolide-lincosamide-streptogramin | 0.067305 | 0.049594 | 0.048992 | 0.051172 | 0.017844 | 0.075317 |
| multidrug | 0.006096 | 0.011224 | 0.003433 | 0.004914 | 0.005123 | 0.004877 |
| polymyxin | 1.56E-05 | 0 | 2.21E-05 | 3.44E-05 | 7.06E-06 | 5.68E-05 |
| puromycin | 0 | 0 | 0 | 0 | 0 | 0 |
| quinolone | 0 | 0 | 0 | 0 | 0 | 0 |
| rifamycin | 0 | 0 | 0 | 0 | 2.43E-05 | 0 |
| spectinomycin | 0 | 0 | 0 | 0 | 0 | 0 |
| sulfonamide | 0 | 0 | 0 | 0 | 0 | 0.000322 |
| tetracenomycin_C | 0 | 0 | 0 | 0 | 0 | 0 |
| tetracycline | 0.125803 | 0.112519 | 0.144419 | 0.142764 | 0.169773 | 0.158462 |
| trimethoprim | 0 | 0 | 0 | 0 | 0 | 0 |
| unclassified | 0.000275 | 0.00033 | 0.000325 | 0.000519 | 0.000302 | 0.000478 |
| vancomycin | 0.011594 | 0.012279 | 0.026468 | 0.031164 | 0.04041 | 0.061922 |
| SUM | 0.363455 | 0.287874 | 0.33126 | 0.515311 | 0.446267 | 0.367376 |

| ARG_types | 8014_8W | 8014_10W |
|---|---|---|
| aminoglycoside | 0.056465 | 0.026263 |
| bacitracin | 0.012715 | 0.030806 |
| beta-lactam | 0.06502 | 0.028653 |
| bleomycin | 0 | 0 |
| carbomycin | 0 | 0 |
| chloramphenicol | 0.000895 | 0.000664 |
| fosfomycin | 0 | 0 |
| fosmidomycin | 0.001562 | 0.001177 |
| fusaric-acid | 0 | 0 |
| fusidic-acid | 0 | 0 |
| kasugamycin | 0 | 0.000339 |
| macrolide-lincosamide-streptogramin | 0.077282 | 0.089888 |
| multidrug | 0.005789 | 0.013432 |
| polymyxin | 2.48E-05 | 0.000496 |
| puromycin | 0 | 0 |
| quinolone | 1.51E-05 | 8.48E-06 |
| rifamycin | 1.51E-05 | 0 |
| spectinomycin | 0 | 0 |
| sulfonamide | 0.000662 | 4.01E-05 |
| tetracenomycin_C | 0 | 0 |
| tetracycline | 0.169782 | 0.154553 |
| trimethoprim | 0 | 2.20E-05 |
| unclassified | 0.000391 | 0.002167 |
| vancomycin | 0.02999 | 0.06274 |
| SUM | 0.420609 | 0.411247 |
