## Supplemental Table 10 resistome for "High prevalence of antibiotic resistance in traditionally fermented foods as a critical risk factor for host gut antibiotic resistome"

Supplemental Table 10. Fecal antibiotic resistome of subjects by diets high in plant fibers.

| ARG_types | 8009_B2W | 8009_0W | 8009_8W | 8009_10W | 8041_B2W | 8041_0W |
|---|---|---|---|---|---|---|
| aminoglycoside | 0.017951 | 0.045349 | 0.029031 | 0.03716 | 0.034937 | 0.043865 |
| bacitracin | 0.014153 | 0.024068 | 0.02261 | 0.040447 | 0.020147 | 0.017826 |
| beta-lactam | 0.012283 | 0.048176 | 0.054454 | 0.049828 | 0.078317 | 0.05107 |
| bleomycin | 0 | 0 | 0 | 0 | 0 | 0 |
| carbomycin | 0 | 0 | 0 | 0 | 0 | 0 |
| chloramphenicol | 0.00138 | 0.002333 | 0.001527 | 0.001027 | 0.00156 | 0.000455 |
| fosfomycin | 0 | 0 | 0 | 0 | 0.00E+00 | 0 |
| fosmidomycin | 0.000932 | 0.002235 | 0.00195 | 0.001486 | 0.002678 | 0.002967 |
| fusaric-acid | 0 | 0 | 0 | 0 | 0 | 0 |
| fusidic-acid | 0 | 0 | 0 | 0 | 0 | 0 |
| kasugamycin | 0 | 0.000185 | 0.000355 | 2.30E-05 | 0.00E+00 | 0 |
| macrolide-lincosamide-streptogramin | 0.027864 | 0.059653 | 0.069701 | 0.064788 | 0.130387 | 0.112107 |
| multidrug | 0.002959 | 0.004994 | 0.005897 | 0.004992 | 0.002662 | 0.00921 |
| polymyxin | 1.84E-05 | 2.90E-05 | 6.78E-05 | 9.49E-06 | 0.00E+00 | 0 |
| puromycin | 0 | 0 | 0 | 0 | 0 | 0 |
| quinolone | 0 | 0 | 0 | 0 | 0 | 0.000143 |
| rifamycin | 0 | 0 | 0 | 0 | 0.00E+00 | 7.51E-05 |
| spectinomycin | 0 | 0 | 0 | 0 | 0 | 0 |
| sulfonamide | 0.001146 | 0.00199 | 0.002458 | 0.001363 | 0.006115 | 0.003042 |
| tetracenomycin_C | 0 | 0 | 0 | 0 | 0 | 0 |
| tetracycline | 0.093841 | 0.240322 | 0.208632 | 0.211553 | 0.199122 | 0.22006 |
| trimethoprim | 0 | 0 | 0 | 0 | 0 | 9.58E-05 |
| unclassified | 0.000277 | 0.000557 | 0.00119 | 0.000315 | 0.000139 | 0.000724 |
| vancomycin | 0.009957 | 0.019526 | 0.017214 | 0.020881 | 0.023564 | 0.040059 |
| SUM | 0.182759 | 0.449417 | 0.415087 | 0.433871 | 0.499627 | 0.501698 |

| ARG_types | 8041_8W | 8041_10W | 8038_B2W | 8038_0W | 8038_8W | 8038_10W |
|---|---|---|---|---|---|---|
| aminoglycoside | 0.027263 | 0.033049 | 0.052453 | 0.016903 | 0.036582 | 0.036787 |
| bacitracin | 0.015201 | 0.016654 | 0.011498 | 0.02088 | 0.022867 | 0.018901 |
| beta-lactam | 0.083315 | 0.067393 | 0.02146 | 0.009822 | 0.020975 | 0.019126 |
| bleomycin | 0 | 0 | 0 | 0 | 0 | 0 |
| carbomycin | 0 | 0 | 0 | 0 | 0 | 0 |
| chloramphenicol | 0 | 0.000512 | 0.001047 | 0.000461 | 0.000587 | 0.000595 |
| fosfomycin | 0 | 0 | 0 | 0 | 0 | 0 |
| fosmidomycin | 0.004026 | 0.002464 | 0.003717 | 0.002361 | 0.003405 | 0.002103 |
| fusaric-acid | 0 | 0 | 0 | 0 | 0 | 0 |
| fusidic-acid | 0 | 0 | 0 | 0 | 0 | 0 |
| kasugamycin | 4.65E-04 | 0 | 7.39E-05 | 0 | 0.000308 | 0 |
| macrolide-lincosamide-streptogramin | 0.132589 | 0.135412 | 0.070872 | 0.040765 | 0.065668 | 0.052661 |
| multidrug | 0.015924 | 0.003406 | 0.003126 | 0.005005 | 0.006826 | 0.003223 |
| polymyxin | 0.000499 | 0 | 0.000165 | 0.000182 | 0.000147 | 0.000127 |
| puromycin | 0 | 0 | 0 | 0 | 0 | 0 |
| quinolone | 0.004269 | 0 | 0 | 0 | 0 | 0 |
| rifamycin | 0 | 0.00E+00 | 0 | 0 | 0 | 0.00E+00 |
| spectinomycin | 0 | 0 | 0 | 0 | 0 | 0 |
| sulfonamide | 0.003977 | 0.013617 | 0.000882 | 9.77E-04 | 0.000435 | 0.000692 |
| tetracenomycin_C | 0 | 0 | 0 | 0 | 0 | 0 |
| tetracycline | 0.190779 | 0.23013 | 0.163941 | 0.160417 | 0.24305 | 0.16526 |
| trimethoprim | 0.001505 | 0 | 0 | 0 | 0 | 0 |
| unclassified | 0.003708 | 0.000353 | 0.000157 | 0.000324 | 0.001441 | 0.000141 |
| vancomycin | 0.027037 | 0.023238 | 0.009121 | 0.011369 | 0.018685 | 0.011512 |
| SUM | 0.510558 | 0.526228 | 0.338513 | 0.269464 | 0.420975 | 0.311128 |

| ARG_types | 8017_B2W | 8017_0W | 8017_10W | 8037_B2W | 8037_0W | 8037_8W |
|---|---|---|---|---|---|---|
| aminoglycoside | 0.0167 | 0.007699 | 0.00971 | 0.052423 | 0.028569 | 0.038102 |
| bacitracin | 0.042928 | 0.022556 | 0.029687 | 0.020654 | 0.021097 | 0.022058 |
| beta-lactam | 0.045067 | 0.049148 | 0.080903 | 0.040738 | 0.028462 | 0.026846 |
| bleomycin | 0 | 0 | 0 | 0 | 0 | 0 |
| carbomycin | 0 | 0 | 0 | 0 | 0 | 0 |
| chloramphenicol | 0.006355 | 0.001437 | 0.002573 | 0.001039 | 0.001395 | 0.0007 |
| fosfomycin | 0 | 0 | 0 | 0 | 0 | 0 |
| fosmidomycin | 0.013636 | 0.000927 | 0.000852 | 0.00043 | 0.001539 | 0.000662 |
| fusaric-acid | 0 | 0 | 0 | 0 | 0 | 0 |
| fusidic-acid | 0 | 0 | 0 | 0 | 0 | 0 |
| kasugamycin | 0.01365 | 0.00031 | 7.17E-05 | 3.21E-05 | 0.000374 | 0.000105 |
| macrolide-lincosamide-streptogramin | 0.061069 | 0.022559 | 0.066639 | 0.053121 | 0.034061 | 0.038649 |
| multidrug | 0.33417 | 0.013823 | 0.006444 | 0.005906 | 0.011427 | 0.00605 |
| polymyxin | 0.011983 | 0.000345 | 0.000341 | 1.25E-04 | 0.000194 | 0.00E+00 |
| puromycin | 0 | 0 | 0 | 0 | 0 | 0 |
| quinolone | 0.001375 | 0.003071 | 6.61E-05 | 0 | 0 | 0 |
| rifamycin | 0 | 0 | 0 | 0 | 0 | 0 |
| spectinomycin | 0 | 0 | 0 | 0 | 0 | 0 |
| sulfonamide | 0.014285 | 0.001376 | 0 | 0 | 0 | 0 |
| tetracenomycin_C | 0 | 0 | 0 | 0 | 0 | 0 |
| tetracycline | 0.100834 | 0.098916 | 0.125105 | 0.168911 | 0.177608 | 0.13829 |
| trimethoprim | 0.004021 | 0.000383 | 0 | 0 | 0 | 0.00E+00 |
| unclassified | 0.078715 | 0.002116 | 0.000656 | 0.000857 | 0.002463 | 0.001421 |
| vancomycin | 0.0206 | 0.022079 | 0.019944 | 0.010403 | 0.012974 | 0.010478 |
| SUM | 0.765387 | 0.246743 | 0.342993 | 0.354639 | 0.320164 | 0.283361 |

| ARG_types | 8037_10W | 8036_B2W | 8036_0W | 8036_10W | 8029_B2W | 8029_0W |
| --- | --- | --- | --- | --- | --- | --- |
| aminoglycoside | 0.047054 | 0.011905 | 0.015572 | 0.003533 | 0.051956 | 0.030459 |
| bacitracin | 0.021635 | 0.022238 | 0.014588 | 0.018714 | 0.032348 | 0.016111 |
| beta-lactam | 0.048199 | 0.034649 | 0.054258 | 0.069753 | 0.031508 | 0.034856 |
| bleomycin | 0 | 0 | 0 | 0 | 0 | 0 |
| carbomycin | 0 | 0 | 0 | 0 | 0 | 0 |
| chloramphenicol | 0.000778 | 0.000182 | 0.000341 | 0 | 8.77E-03 | 0.002617 |
| fosfomycin | 0 | 0 | 0.000251 | 0 | 0 | 0.00E+00 |
| fosmidomycin | 0.001344 | 0.001228 | 0.001531 | 0.002604 | 0.002077 | 0.001672 |
| fusaric-acid | 0 | 0 | 0 | 0 | 0 | 0 |
| fusidic-acid | 0 | 0 | 0 | 0 | 0 | 0 |
| kasugamycin | 4.55E-05 | 4.42E-05 | 4.13E-05 | 0 | 3.95E-05 | 0.000104 |
| macrolide-lincosamide-streptogramin | 0.059769 | 0.056465 | 0.073207 | 0.078655 | 0.042873 | 0.026733 |
| multidrug | 0.006572 | 0.004791 | 0.004497 | 0.003512 | 0.009044 | 0.005051 |
| polymyxin | 2.05E-05 | 8.13E-05 | 0.000101 | 0.00E+00 | 5.96E-06 | 2.52E-05 |
| puromycin | 0 | 0 | 0 | 0 | 0 | 0 |
| quinolone | 0 | 0.000271 | 3.92E-05 | 1.95E-05 | 0 | 0 |
| rifamycin | 0.00E+00 | 0 | 0 | 0 | 0 | 0 |
| spectinomycin | 0 | 0 | 0 | 0 | 0 | 0 |
| sulfonamide | 0 | 3.44E-05 | 7.2E-05 | 7.12E-05 | 3.9E-05 | 0 |
| tetracenomycin_C | 0 | 0 | 0 | 0 | 0 | 0 |
| tetracycline | 0.189239 | 0.235827 | 0.241631 | 0.262887 | 0.201681 | 0.177969 |
| trimethoprim | 0 | 9.34E-05 | 0 | 0 | 3.3E-05 | 0 |
| unclassified | 0.001392 | 0.000661 | 0.001022 | 0.000286 | 0.000735 | 0.000892 |
| vancomycin | 0.008607 | 0.013158 | 0.014574 | 0.013895 | 0.016546 | 0.016837 |
| SUM | 0.384654 | 0.381629 | 0.421726 | 0.45393 | 0.397652 | 0.313326 |

| ARG_types | 8029_8W | 8029_10W | 8002_B2W | 8002_0W | 8002_8W | 8002_10W |
|---|---|---|---|---|---|---|
| aminoglycoside | 0.026638 | 0.024156 | 0.011026 | 0.00634 | 0.010514 | 0.007812 |
| bacitracin | 0.024728 | 0.01843 | 0.012955 | 0.035643 | 0.01843 | 0.047663 |
| beta-lactam | 0.042315 | 0.030006 | 0.033407 | 0.033729 | 0.02913 | 0.023528 |
| bleomycin | 0 | 0 | 0 | 0 | 0 | 0 |
| carbomycin | 0 | 0 | 0 | 0 | 0 | 0 |
| chloramphenicol | 0.008467 | 0.011809 | 0.000387 | 0.00026 | 0.000142 | 0.000386 |
| fosfomycin | 0 | 0 | 0 | 0.00E+00 | 0 | 0 |
| fosmidomycin | 0.001921 | 0.002029 | 0.000825 | 0.00051 | 0.00101 | 0.001161 |
| fusaric-acid | 0 | 0 | 0 | 0 | 0 | 0 |
| fusidic-acid | 0 | 0 | 0 | 0 | 0 | 0 |
| kasugamycin | 1.45E-05 | 8.17E-05 | 0.000255 | 0 | 2.65E-05 | 0 |
| macrolide-lincosamide-streptogramin | 0.023625 | 0.022783 | 0.028509 | 0.025388 | 0.02444 | 0.018549 |
| multidrug | 0.004026 | 0.007611 | 0.013215 | 0.004796 | 0.004722 | 0.004221 |
| polymyxin | 0 | 8.68E-05 | 0.000442 | 0 | 4.39E-05 | 0 |
| puromycin | 0 | 0 | 0 | 0 | 0 | 0 |
| quinolone | 0 | 0.00E+00 | 1.03E-05 | 0 | 0 | 0 |
| rifamycin | 0 | 0 | 0 | 0 | 0 | 0 |
| spectinomycin | 0 | 0 | 0 | 0 | 0 | 0 |
| sulfonamide | 0 | 0 | 0.00071 | 0 | 0 | 0 |
| tetracenomycin_C | 0 | 0 | 0 | 0 | 0 | 0 |
| tetracycline | 0.177067 | 0.15381 | 0.160623 | 0.161081 | 0.163504 | 0.139511 |
| trimethoprim | 0 | 0 | 0 | 0 | 0 | 0 |
| unclassified | 0.00014 | 0.000471 | 0.003177 | 0.000988 | 0.000622 | 0.000305 |
| vancomycin | 0.017835 | 0.010681 | 0.008667 | 0.020912 | 0.011908 | 0.022386 |
| SUM | 0.326775 | 0.281955 | 0.274209 | 0.289648 | 0.264493 | 0.265522 |

| ARG_types | 8003_B2W | 8003_0W | 8003_8W | 8003_10W | 8039_B2W | 8039_0W |
|---|---|---|---|---|---|---|
| aminoglycoside | 0.101759 | 0.055531 | 0.056162 | 0.03094 | 0.010795 | 0.018165 |
| bacitracin | 0.013725 | 0.015179 | 0.012235 | 0.013799 | 0.029236 | 0.024801 |
| beta-lactam | 0.033142 | 0.073877 | 0.082018 | 0.140024 | 0.050688 | 0.090773 |
| bleomycin | 0 | 0 | 0 | 0 | 0 | 0 |
| carbomycin | 0 | 0 | 0.00E+00 | 0 | 0 | 0 |
| chloramphenicol | 0.009381 | 0.00568 | 0.006642 | 0.00311 | 0.000157 | 0.0009 |
| fosfomycin | 0.003238 | 0.001607 | 0.002193 | 0.001179 | 0 | 0 |
| fosmidomycin | 0.001249 | 0.000885 | 0.001384 | 0.001493 | 0.000485 | 0.001385 |
| fusaric-acid | 0 | 0 | 0 | 0 | 0 | 0 |
| fusidic-acid | 0 | 0 | 0 | 0 | 0 | 0 |
| kasugamycin | 0.000181 | 0 | 1.60E-04 | 0.000373 | 0.000109 | 0.000462 |
| macrolide-lincosamide-streptogramin | 0.073534 | 0.054946 | 0.044122 | 0.029885 | 0.07748 | 0.125935 |
| multidrug | 0.006072 | 0.004343 | 0.004129 | 0.003103 | 0.007786 | 0.022549 |
| polymyxin | 1.67E-04 | 0.000169 | 0.000114 | 1.71E-05 | 7.27E-05 | 0.000733 |
| puromycin | 0 | 0 | 0 | 0 | 0 | 0 |
| quinolone | 0 | 0 | 0 | 0 | 0 | 0 |
| rifamycin | 0.00E+00 | 0 | 0 | 0 | 0 | 0 |
| spectinomycin | 0 | 0 | 0 | 0 | 0 | 0 |
| sulfonamide | 0.000177 | 0.000765 | 6.06E-04 | 2.72E-04 | 0 | 0 |
| tetracenomycin_C | 0 | 0 | 0 | 0 | 0 | 0 |
| tetracycline | 0.323506 | 0.212959 | 0.186425 | 0.151616 | 0.218654 | 0.251607 |
| trimethoprim | 0 | 0 | 0 | 0 | 0 | 0 |
| unclassified | 0.000462 | 0.000502 | 0.000654 | 0.001153 | 0.00088 | 0.005159 |
| vancomycin | 0.057469 | 0.054865 | 0.050691 | 0.018024 | 0.013125 | 0.009815 |
| SUM | 0.624062 | 0.481308 | 0.447535 | 0.394988 | 0.409468 | 0.552285 |

| ARG_types | 8039_8W | 8039_10W | 8006_B2W | 8006_0W | 8006_8W | 8006_10W |
|---|---|---|---|---|---|---|
| aminoglycoside | 0.008036 | 0.014238 | 0.011363 | 0.007024 | 0.019454 | 0.011582 |
| bacitracin | 0.03393 | 0.02391 | 0.020021 | 0.01814 | 0.025746 | 0.025591 |
| beta-lactam | 0.107531 | 0.126718 | 0.013801 | 0.018815 | 0.004441 | 0.017358 |
| bleomycin | 0 | 0 | 0 | 0 | 0 | 0 |
| carbomycin | 0 | 0 | 0 | 0 | 0 | 0 |
| chloramphenicol | 0.000251 | 0.000514 | 0.003994 | 0.002253 | 0.014463 | 0.004912 |
| fosfomycin | 0.00E+00 | 0.00E+00 | 0 | 0 | 0 | 0 |
| fosmidomycin | 0.001244 | 0.000545 | 0.002277 | 0.003118 | 0.001142 | 0.002683 |
| fusaric-acid | 0 | 0 | 0 | 0 | 0 | 0 |
| fusidic-acid | 0 | 0 | 0 | 0 | 0 | 0 |
| kasugamycin | 0.000232 | 0 | 7.75E-05 | 0 | 8.05E-05 | 0.000102 |
| macrolide-lincosamide-streptogramin | 0.125023 | 0.132023 | 0.004383 | 0.005591 | 0.00459 | 0.010953 |
| multidrug | 0.013 | 0.002176 | 0.003995 | 0.002877 | 0.004324 | 0.004463 |
| polymyxin | 0.000277 | 0 | 6.44E-05 | 2.41E-05 | 4.81E-05 | 0.000225 |
| puromycin | 0 | 0 | 0 | 0 | 0 | 0 |
| quinolone | 0 | 0 | 9.93E-06 | 7.76E-05 | 0.000448 | 0.0001 |
| rifamycin | 0 | 0 | 0 | 0 | 0 | 0 |
| spectinomycin | 0 | 0 | 0 | 0 | 0 | 0 |
| sulfonamide | 0 | 0 | 0.000247 | 3.42E-05 | 0 | 9.06E-05 |
| tetracenomycin_C | 0 | 0 | 0 | 0 | 0 | 0 |
| tetracycline | 0.249684 | 0.247437 | 0.139534 | 0.147967 | 0.129128 | 0.152511 |
| trimethoprim | 0 | 0 | 4.1E-05 | 0 | 4.78E-05 | 0 |
| unclassified | 0.002231 | 0.000238 | 0.000879 | 0.000276 | 0.000475 | 0.001027 |
| vancomycin | 0.009715 | 0.007343 | 0.018546 | 0.013768 | 0.029569 | 0.021426 |
| SUM | 0.551152 | 0.555141 | 0.219233 | 0.219967 | 0.233956 | 0.253023 |

| ARG_types | 8001_0W | 8001_8W | 8001_10W | 8023_0W | 8023_8W | 8023_10W |
|---|---|---|---|---|---|---|
| aminoglycoside | 0.021815 | 0.030157 | 0.026864 | 0.008388 | 0.018973 | 0.007669 |
| bacitracin | 0.020398 | 0.025485 | 0.029305 | 0.01808 | 0.023963 | 0.021696 |
| beta-lactam | 0.076979 | 0.037186 | 0.030727 | 0.12621 | 0.061971 | 0.148234 |
| bleomycin | 0 | 0 | 0 | 0 | 0 | 0 |
| carbomycin | 0 | 0 | 0 | 0 | 0 | 0 |
| chloramphenicol | 0.003115 | 0.001923 | 0.00198 | 0.000448 | 0.000366 | 5.27E-05 |
| fosfomycin | 0 | 0 | 0 | 0 | 0 | 0 |
| fosmidomycin | 7.66E-04 | 2.24E-03 | 1.06E-03 | 0.001389 | 0.001057 | 0.001378 |
| fusaric-acid | 0 | 0 | 0 | 0 | 0 | 0 |
| fusidic-acid | 0 | 0 | 0 | 0 | 0 | 0 |
| kasugamycin | 0 | 0 | 1.60E-04 | 0.000182 | 0.000203 | 0.000516 |
| macrolide-lincosamide-streptogramin | 0.044885 | 0.02781 | 0.040247 | 0.021998 | 0.053068 | 0.022141 |
| multidrug | 0.008895 | 0.004608 | 0.005629 | 0.007336 | 0.004115 | 0.00332 |
| polymyxin | 2.78E-05 | 4.20E-05 | 3.07E-05 | 2.32E-04 | 0.00E+00 | 9.77E-05 |
| puromycin | 0 | 0 | 0 | 0 | 0 | 0 |
| quinolone | 0.000245 | 0.000308 | 0.00021 | 0 | 0.00E+00 | 0.00E+00 |
| rifamycin | 0 | 0 | 0.00E+00 | 0 | 0.00E+00 | 0 |
| spectinomycin | 0 | 0 | 0 | 0 | 0 | 0 |
| sulfonamide | 0 | 0.000178 | 0.000411 | 0.000458 | 1.74E-05 | 4.36E-05 |
| tetracenomycin_C | 0 | 0 | 0 | 0 | 0 | 0 |
| tetracycline | 0.168752 | 0.152184 | 0.137074 | 0.267121 | 0.194696 | 0.285508 |
| trimethoprim | 0 | 0 | 0 | 0.000223 | 0 | 7.50E-05 |
| unclassified | 0.000705 | 0.00065 | 0.000768 | 0.001515 | 0.000646 | 0.001441 |
| vancomycin | 0.029834 | 0.033258 | 0.035701 | 0.014157 | 0.02651 | 0.015128 |
| SUM | 0.376419 | 0.316027 | 0.310167 | 0.467738 | 0.385584 | 0.507301 |

| ARG_types | 8018_B2W | 80186_8W | 8018_10W | 8022_B2W | 8022_8W | 8022_10W |
|---|---|---|---|---|---|---|
| aminoglycoside | 0.005082 | 0.000795 | 0.000714 | 0.018066 | 0.053259 | 0.039421 |
| bacitracin | 0.019425 | 0.018306 | 0.0186 | 0.009919 | 0.020146 | 0.014617 |
| beta-lactam | 0.032357 | 0.062896 | 0.084634 | 0.02637 | 0.029169 | 0.031086 |
| bleomycin | 0 | 0 | 0 | 0 | 0 | 0 |
| carbomycin | 0 | 0 | 0 | 0 | 0 | 0 |
| chloramphenicol | 0.001032 | 0.002663 | 0.003006 | 0.000108 | 0.000583 | 0.000316 |
| fosfomycin | 0 | 0 | 0 | 0 | 0 | 0 |
| fosmidomycin | 0.000727 | 0.000482 | 0.000631 | 0.002393 | 0.005965 | 0.00343 |
| fusaric-acid | 0 | 0 | 0 | 0 | 0 | 0 |
| fusidic-acid | 0 | 0 | 0 | 0 | 0 | 0 |
| kasugamycin | 0.000232 | 3.48E-05 | 3.50E-05 | 0.001523 | 0.004161 | 0.003272 |
| macrolide-lincosamide-streptogramin | 0.023743 | 0.022435 | 0.016271 | 0.029797 | 0.037001 | 0.040499 |
| multidrug | 0.010579 | 0.002537 | 0.003363 | 0.0341 | 0.117701 | 0.071975 |
| polymyxin | 0.000378 | 0 | 2.26E-05 | 0.001304 | 0.003624 | 0.002506 |
| puromycin | 0 | 0 | 0 | 0 | 0 | 0 |
| quinolone | 3.71E-05 | 0.000308 | 0.00062 | 0 | 0 | 0 |
| rifamycin | 0 | 0 | 0 | 0 | 0 | 0 |
| spectinomycin | 0 | 0 | 0 | 0 | 0 | 0 |
| sulfonamide | 0.000404 | 0.000154 | 7.36E-05 | 0.00329 | 0.007895 | 0.00533 |
| tetracenomycin_C | 0 | 0 | 0 | 0 | 0 | 0 |
| tetracycline | 0.148497 | 0.12318 | 0.142695 | 0.211351 | 0.199801 | 0.195577 |
| trimethoprim | 0 | 0 | 0 | 0.001317 | 0.006573 | 0.003831 |
| unclassified | 0.00232 | 0.00065 | 0.000886 | 0.009418 | 0.026291 | 0.017703 |
| vancomycin | 0.014968 | 0.007357 | 0.008329 | 0.016018 | 0.022383 | 0.018487 |
| SUM | 0.259782 | 0.241797 | 0.279882 | 0.364975 | 0.534551 | 0.44805 |
